# Diversity in the impact of heterogeneities on recurrent networks performing a cognitive task

**DOI:** 10.1101/2025.01.20.633872

**Authors:** Anjana Santhosh, Rishikesh Narayanan

**Affiliations:** Cellular Neurophysiology Laboratory, Molecular Biophysics Unit, Indian Institute of Science, Bangalore, India

**Keywords:** recurrent network, cognitive task, heterogeneity, variability, hyperparameters, robustness

## Abstract

**Background and motivation:** Artificial recurrent networks are widely used as models to study the complex dynamics underlying biological neural networks during execution of cognitive tasks. However, most studies assume individual units in the recurrent network to be homogeneous repeating units, whereas real neurons exhibit several forms of heterogeneities. In this study, we designed and employed a systematic framework for quantitative assessment of the impact of neural heterogeneities on recurrent networks that were trained to perform a cognitive task.

**Methodology:** Our framework involved training of a population of recurrent networks, differing in terms of their hyperparameters, to perform a cognitive task in the presence of six graded levels of intrinsic heterogeneities. We tested the impact of heterogeneities on several performance metrics that encompassed training performance, task-execution dynamics, and resilience to different forms of post-training heterogeneities (also introduced at different levels).

**Results:** Our population-of-networks approach demonstrate that intrinsic heterogeneities impacted network performance and dynamics in diverse ways even if they were trained with the same training algorithm, convergence criteria, and task specifications. First, our analyses unveiled pronounced network-to-network variability in the dependence of training performance on the level of heterogeneity, in terms of the number of training trials required for learning and the error values associated with task performance. Second, the impact of training heterogeneities on network dynamics during task execution also manifested substantial variability across networks. Finally, our analyses revealed a prominent impact of different forms of post-training heterogeneities on performance errors and network dynamics. We observed progressive increases in errors as well as in trajectory deviations with graded increases in post-training heterogeneities. Importantly, we observed pronounced variability in how robustness to post-training heterogeneities depended on the level of training heterogeneities. Specifically, certain networks showed enhanced robustness to post-training heterogeneities when training heterogeneities were low, whereas others showed better robustness when training heterogeneities were high.

**Implications:** The striking nature of network-to-network variability observed in our analyses strongly advocates a complex systems viewpoint to study the impact of neural heterogeneities on circuit function. Within such a complex system framework, where several functionally specialized subsystems interact with each other in non-random ways to yield collective performance of the task, we argue that the emphasis should not be on heterogeneities in *individual* components of neural circuits. Instead, we emphasize the need to focus on the *global structure* of different forms and degrees of heterogeneities across different components and a systematic assessment of how they *interact* with each other towards *adapting* and achieving collective function.

## INTRODUCTION

Artificial recurrent networks are widely used as models to study the complex dynamics underlying biological neural networks during execution of cognitive tasks (Sussillo and Abbott, 2009; Mante et al., 2013; Barak, 2017; Yang et al., 2019; Gallego et al., 2020; Safaie et al., 2023; Driscoll et al., 2024). However, most studies assume individual units in the recurrent network to be homogeneous repeating units, an assumption that is fundamentally in conflict with biological neural networks where individual neurons manifest several forms of heterogeneities (Marder, 2011; Cadwell et al., 2016; Cembrowski and Spruston, 2019; Mishra and Narayanan, 2020; Huckleberry and Shansky, 2021; Nandi et al., 2022; Han et al., 2023; Langlieb et al., 2023; Piwecka et al., 2023; Siletti et al., 2023; Dembrow et al., 2024). There are several lines of evidence that these heterogeneities are not just chance occurrences but have diverse functional impact on different aspects of neural and network function. A diversity of such impact has been reported in terms of providing alternate routes to implement specific functional outcomes (Prinz et al., 2004; Cropper et al., 2016; Mishra and Narayanan, 2019; Rathour and Narayanan, 2019; Wang et al., 2019; Albantakis et al., 2024; Mittal and Narayanan, 2024; Calabrese and Marder, 2025), efficient coding (Shamir and Sompolinsky, 2006; Chelaru and Dragoi, 2008; Marsat and Maler, 2010; Ahn et al., 2014; Balasubramanian, 2015; Tesileanu et al., 2019; Seenivasan and Narayanan, 2020; Wada et al., 2021), spatial navigation (Renart et al., 2003; Basak and Narayanan, 2018, 2020; Roy and Narayanan, 2021), decorrelation (Padmanabhan and Urban, 2010; Tripathy et al., 2013; Mishra and Narayanan, 2019), disruption of function in certain types of networks (Mittal and Narayanan, 2021; Gast et al., 2024), introducing plasticity heterogeneity towards efficient resource allocation (Yiu et al., 2014; Park et al., 2016; Josselyn and Frankland, 2018; Rao-Ruiz et al., 2019; Lau et al., 2020; Shridhar et al., 2022), modulating synchrony in networks (Wang and Buzsaki, 1996; White et al., 1998; Brunel and Hakim, 1999; Maex and De Schutter, 2003; Tikidji-Hamburyan et al., 2015; Gast et al., 2024), promoting stability of learning in spiking neural networks (Perez-Nieves et al., 2021), and mediating resilience to perturbations (Mishra and Narayanan, 2021a; Ratliff et al., 2021; Gorur-Shandilya et al., 2022; Marder et al., 2022; Rich et al., 2022; Alonso et al., 2023; Marom and Marder, 2023; Schapiro and Marder, 2024; Calabrese and Marder, 2025). Despite the several lines of evidence on the diverse roles of heterogeneities on brain function, the impact of neural heterogeneities on recurrent networks trained to perform a cognitive task has not been explored.

In this study, we develop and employ a framework to systematically assess the role of neural heterogeneities in affecting circuit function. We emphasize five essential features as key to this generic framework to study neural heterogeneities. First, learning in biological systems occur in the presence of heterogeneities and introduces specific structure to the synaptic connectivity and weights (Sussillo and Abbott, 2009; Mante et al., 2013; Barak, 2017; Yang et al., 2019; Gallego et al., 2020; Safaie et al., 2023; Driscoll et al., 2024). Therefore, neural heterogeneities must be introduced during the training process, when the network learns the specific task structure towards fine-tuning its dynamics. Second, as biological heterogeneities could manifest in different forms at different degrees, different forms of heterogeneities must be introduced into the network at multiple levels, specifically designed to assess graded impact of different heterogeneities on performance. Third, a population of networks with very different instantiations of hyperparameters needs to be assessed across all levels of heterogeneities. The use of a single network with a single instantiation of hyperparameters biases outcomes to that choice of hyperparameters, not just in terms of network performance but also in terms of its dependence on heterogeneities. Fourth, biological networks are in a continuous state of flux owing to external perturbations or need to learn additional tasks. It is therefore essential to assess the robustness of network performance to different forms and several levels of post-training heterogeneities. Finally, several quantitative metrics need to be used for evaluating training performance, task-execution dynamics, and resilience to post-training heterogeneities, with the impact of heterogeneities on each of these studied separately across different levels of heterogeneities (both training as well as post-training). Within this broad framework, we explored the impact of intrinsic heterogeneities in recurrent networks performing a cognitive task. We used a modified reward-modulated Hebbian learning mechanism (Miconi, 2017) to train multiple recurrent networks (different hyperparameters), each with several gradations of intrinsic heterogeneities, to perform a simple cognitive task. We quantified training performance, task-execution dynamics, and resilience to different forms of post-training heterogeneities in each of the trained networks towards assessing the impact of training heterogeneities on network performance. Our analyses unveiled pronounced network-to-network variability in the impact of heterogeneities on training, task performance, network dynamics during task execution, and robustness to post-training heterogeneities. Based on our analyses, we propose a complex systems viewpoint to study the impact of neural heterogeneities on circuit function, where the emphasis is not on heterogeneities in individual components. Instead, we emphasize the need to focus on the global structure of different forms and degrees of heterogeneities across different components and how they interact with each other towards achieving collective function.

## RESULTS

The principal goal of this study was to test the impact of neural heterogeneities on a recurrent network trained to perform a cognitive task. We used a fully connected, continuous-time recurrent network with 200 units (Fig. 1). The recurrent network was trained using a modified reward-modulated Hebbian learning algorithm to perform a simple memoryless Go task. Our experimental design towards assessing the impact of heterogeneities on recurrent networks performing this cognitive task involved important methodological constraints. First, we did not binarize heterogeneous *vs.* homogeneous networks. Instead, we introduced heterogeneities in a graded fashion in six different ranges in unit time constants (Fig. 1). Such gradation accounts for variability in the extent of biological heterogeneities observed under different conditions. More importantly, this experimental design involving different levels of heterogeneities (H0–H5) allowed us to assess progressive changes in network performance and dynamics with increasing levels of intrinsic heterogeneities.

**Figure 1.**
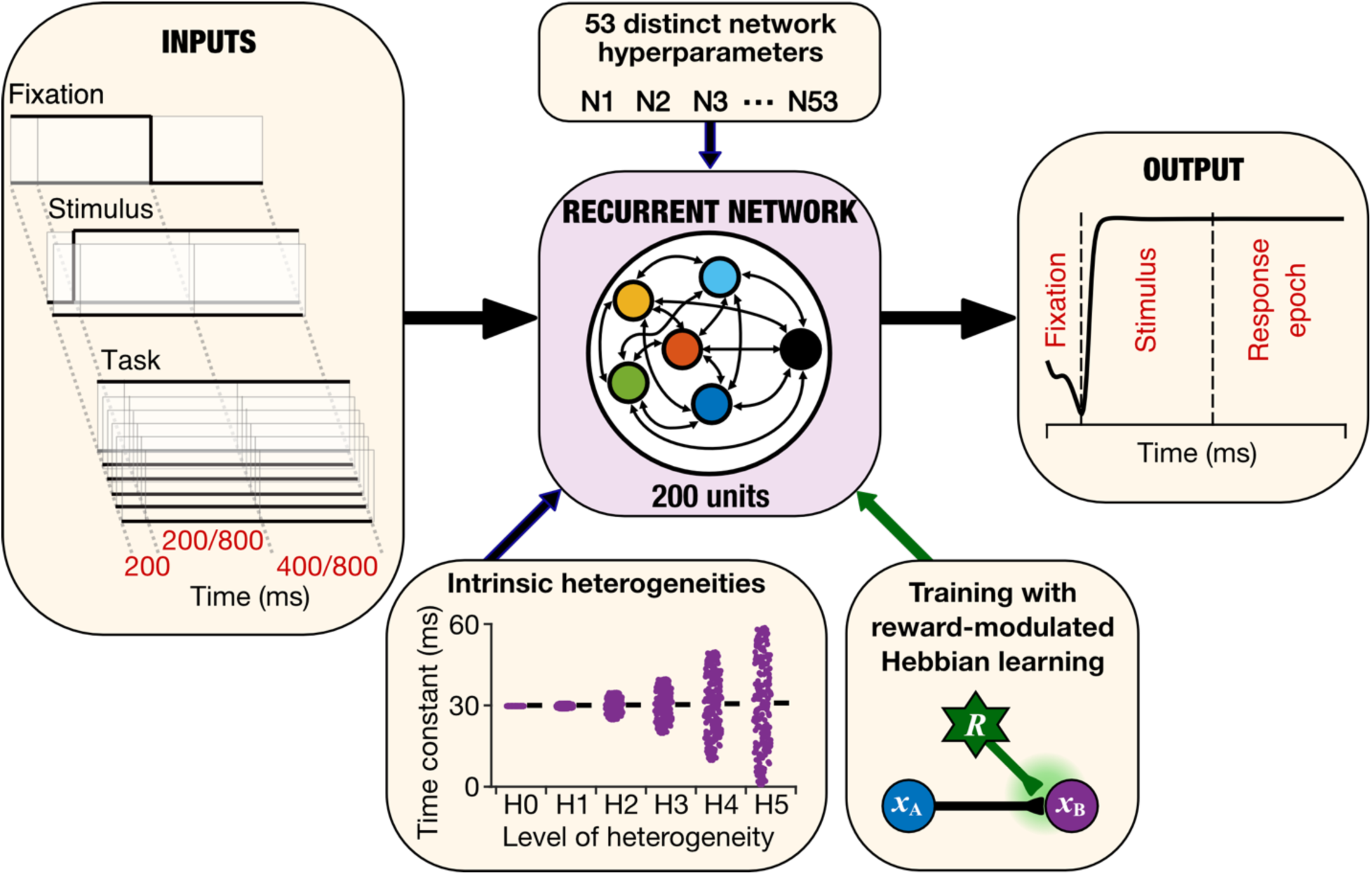
A population of recurrent networks with distinct hyperparameters trained and tested with different levels of heterogeneities to perform a cognitive task. A rate-based fully connected recurrent network of 200 units was initialized with 53 different parametric configurations, each trained and tested with six levels (H0–H5; H0 refers to the homogeneous network) of intrinsic heterogeneities. Total number of networks assessed: 53 × 6 = 318. Progressively higher levels of intrinsic heterogeneities (H1–H5) were achieved by increasing the range of randomly initialized time constants of the units in the network. The network under consideration receives three types of inputs: a fixation input which turns off when the network is expected to provide its output; two inputs corresponding to the stimuli for the task being performed; and 6 distinct inputs which indicate the specific task the network is expected to perform. All networks in this study were trained and tested for the memoryless Go task, which is indicated on the “Inputs” tab. Each trial of the Go task consists of three epochs with variable duration (fixation epoch, fixed at 200 ms; stimulus epoch, switching between 200 or 800 ms; and response epoch, alternating between 400 or 800 ms). A single unit was designated as the output unit of the network whose response was considered as the network’s output. The network was trained using a modified reward-modulated Hebbian learning rule. For the Go task, the expected output is to reflect the input value assigned to either of the two stimuli (which are never simultaneously active).

Second, to avoid biases introduced by the use of a single network for all analyses, we used a population of networks approach throughout our analyses to quantify potential network-to-network variability in training performance and response dynamics. Specifically, we employed 53 instantiations of networks which were different in terms of all hyperparameters that defined initialization of these networks. Together, this yielded 318 different networks (53 different parameter initializations, each with 6 levels of heterogeneity) that were trained individually to perform the cognitive task. The impact of intrinsic heterogeneities on training performance and task execution dynamics was tested for all 53 networks, spanning all 6 levels of heterogeneities.

Third and finally, neural-circuit heterogeneities could be introduced into biological networks after the learning process is complete. As these post-training heterogeneities could also occur with different ranges, our experimental design involved six graded levels each of several forms of post-training heterogeneities (referred to as P0–P5, to distinguish from H0–H5 introduced during training). For each form of post-training heterogeneity, we studied the impact of the 6 levels of post-training heterogeneities on all 318 networks. The experimental design and the associated analyses allowed us to quantitatively assess (i) the extent to which networks were robust to each kind of post-training heterogeneity; (ii) network-to-network variability in how networks react to different forms of post-training heterogeneities; (iii) the impact of pre-training heterogeneities (H0–H5) on how robust the network was to each form of post-training heterogeneity (P0–P5); and (iv) network-to-network variability in the impact of pre-training heterogeneities on network robustness to post-training heterogeneities.

In implementing this broad experimental design, we first trained each of the 318 networks to perform the Go task till each network reached the convergence criterion, whereby the absolute error must be below 0.05 for 100 consecutive training trials. The number of trials required for convergence and the errors across testing trials were noted for each network.

### Pronounced network-to-network variability in the impact of heterogeneities on training performance

All 318 networks successfully reached the set convergence criterion towards executing the Go task. However, the training trajectories manifested pronounced variability across different networks and across heterogeneity levels (Fig. 2*A–C*). First, even in the absence of intrinsic heterogeneities (H0 group), there were large differences in the number of trials required for convergence. Whereas some networks convergence with as low as 4000 trials, others required close to 20000+ trials for achieving convergence (Fig. 2*C*). Second, the impact of progressively increasing the level of heterogeneities (from H0 to H5) increased the number of required training trials in some networks (*e.g.*, N33), decreased in others (*e.g.*, N26), while had no such structure (*e.g.*, N39) in a third set of networks (Fig. 2*A–B*). Similarly, the error values observed in test trials showed pronounced trial-to-trial variability for any given level of heterogeneity (Fig. 2*D*), pronounced network-to-network variability in the error values as well as the dependency on heterogeneities (Fig. 2*D–E*). Specifically, for some networks the median error increased with level of heterogeneity (*e.g.*, N19), for others the median error decreased (*e.g.*, N16), while a third class of networks did not exhibit any monotonic trend (*e.g.*, N18) (Fig. 2*D*).

**Figure 2.**
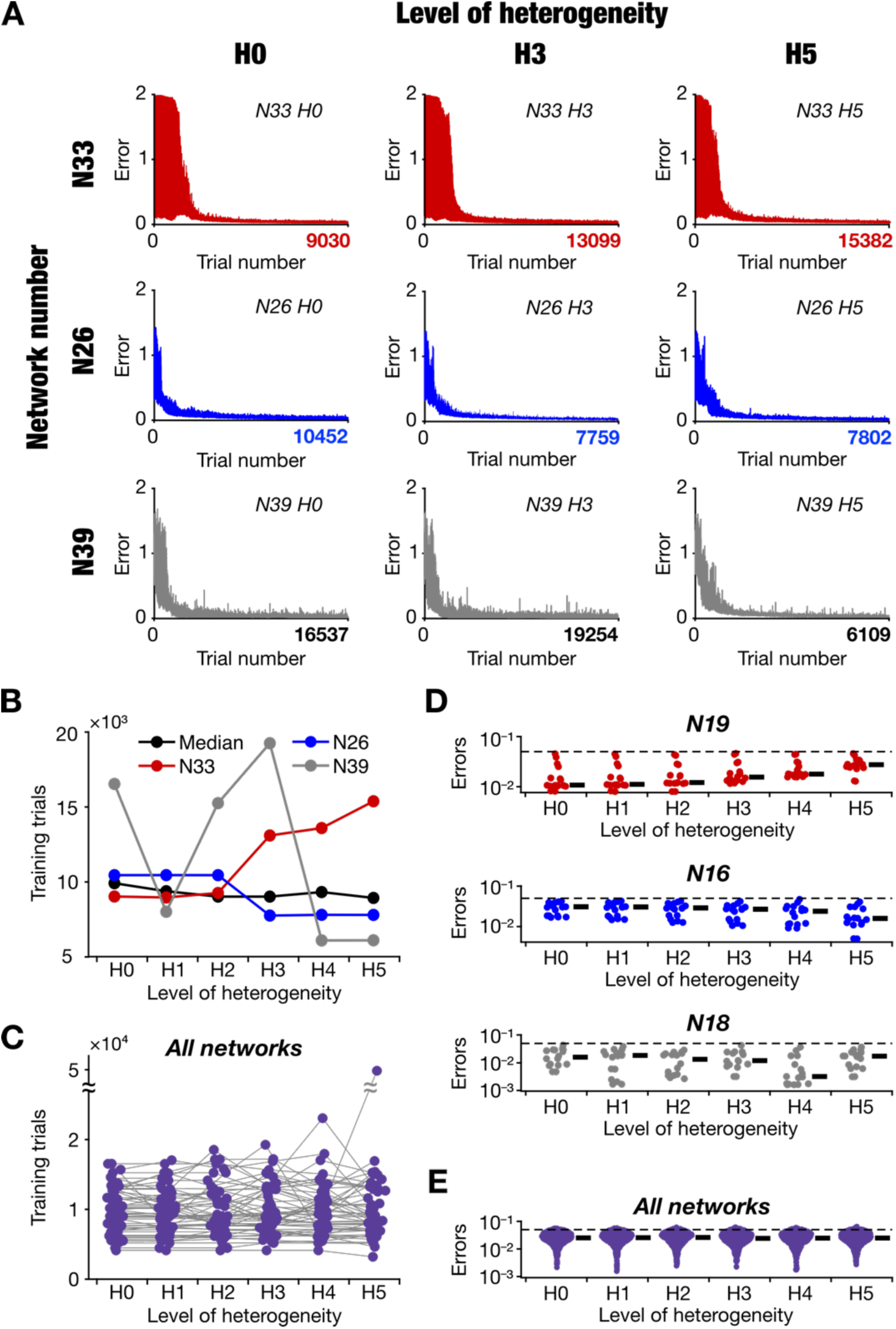
Pronounced network-to-network variability in training performance of heterogeneous recurrent networks trained to perform a cognitive task. *A:* The evolution of absolute error during training for three different initializations (rows) and different levels of intrinsic heterogeneity (columns). The *x*-axis ranges from 0 to 𝑁_!_, the total number of trials required for the specific network to converge. It may be noted that the number of training trials required for convergence increased (N33), decreased (N26), or varied randomly (N39) with increasing levels of heterogeneity. *B:* Number of trials taken to train the three example networks shown in *A* plotted as a function of the level of training heterogeneity, depicting network-to-network variability in the impact of heterogeneities on training. The black line depicts the median number of trials across all 53 networks. *C:* Pronounced network-to-network variability in the number of trials required for convergence in all 53 networks, each trained across the 6 heterogeneity levels (H0–H5). *D:* Distribution of errors in the 16 different testing trials (of the Go task, with different stimuli as well as different durations for the stimulus and the response epochs) plotted across H0–H5 for three example trained networks (dotted lines represent the 0.05 level used as a convergence requirement). Network N19 showed an increase in error with increasing heterogeneity, network N16 manifested a reduction, while network N18 did not show any monotonic pattern. Black dashes indicate median errors for each heterogeneity level. *E:* Distribution of errors in all trials for all 318 trained networks. The black dashes represent the median error for each level of heterogeneity.

For one of the networks (N1), we also tested the impact of intrinsic heterogeneities on networks that were trained with different parameters associated with the network and the training process (Supplementary Fig. S1). The network was trained in the presence of 6 levels of heterogeneities, with a lower learning rate (11 = 0.05), lower connection density (connection density = 50%), or with a more stringent convergence criterion (error below 0.05 for 400 consecutive training trials). With increasing level of heterogeneities, the number of training trials showed an increasing trend in networks trained with a reduced learning rate as well as in the network with a reduced connection density, but the trend reversed in the network that was trained with a stringent convergence criterion (Supplementary Fig. S1*A*). Similarly, we observed variable dependencies of test trial errors on the level of heterogeneities, depending on which parameter was changed in the network (Supplementary Fig. S1*B*). These results demonstrated that the variability in dependence of training performance on the level of heterogeneities extended to other training hyperparameters as well.

Together, these results demonstrated pronounced variability across networks in terms of the number of training trials required for learning (Fig. 2*A–C*), the error values associated with specific trials (Fig. 2*D–E*), connectivity/training hyperparameters (Supplementary Fig. S1), and the dependency of trial numbers (Fig. 2*A–C*) and error values (Fig. 2*D–E*) on the level of intrinsic heterogeneities introduced during training.

### Dynamics of heterogeneous networks during task performance varied across network initializations

We performed principal component analysis (PCA) on the activity dynamics of all recurrent units to visualize and assess the task-associated dynamics of the trained recurrent networks. We repeated this for all 53 networks for each of the 6 different levels of heterogeneities. As the dimensionality of the activity dynamics (equation (10)) was around 3 for this task across all networks, we used a 3-dimensional representation for visualizing the dynamics of these networks (Fig. 3*A*). In this visualization, the initial time point (𝑡 = 0) is represented by a black dot. The trajectory in the reduced dimensional space followed an identical route until the stimulus arrived (Fig. 3*A*). Upon stimulus arrival at the beginning of the stimulus epoch, the trajectory diverged based on the specific input stimuli and eventually converged to one of the four stable points, each defining a specific input pattern (𝒖*_stim_* ∈ {(−1,0), (1,0), (0, −1), (0,1)}). We found the latent space dynamics to be distinct across different levels of heterogeneity for the same network (Supplementary Fig. S2) as well as across different networks for the same level of heterogeneity (Supplementary Fig. S3). To compare trajectories across different networks and across different levels of heterogeneities, we first aligned the latent spaces of the different networks using canonical correlation analysis (CCA) in a pairwise manner using a reference network N1H0 (Fig. 3*A*). Finally, the aligned trajectories were transformed along the principal components of N1H0, thus representing all the network trajectories in the same latent space (Fig 3*A*).

**Figure 3.**
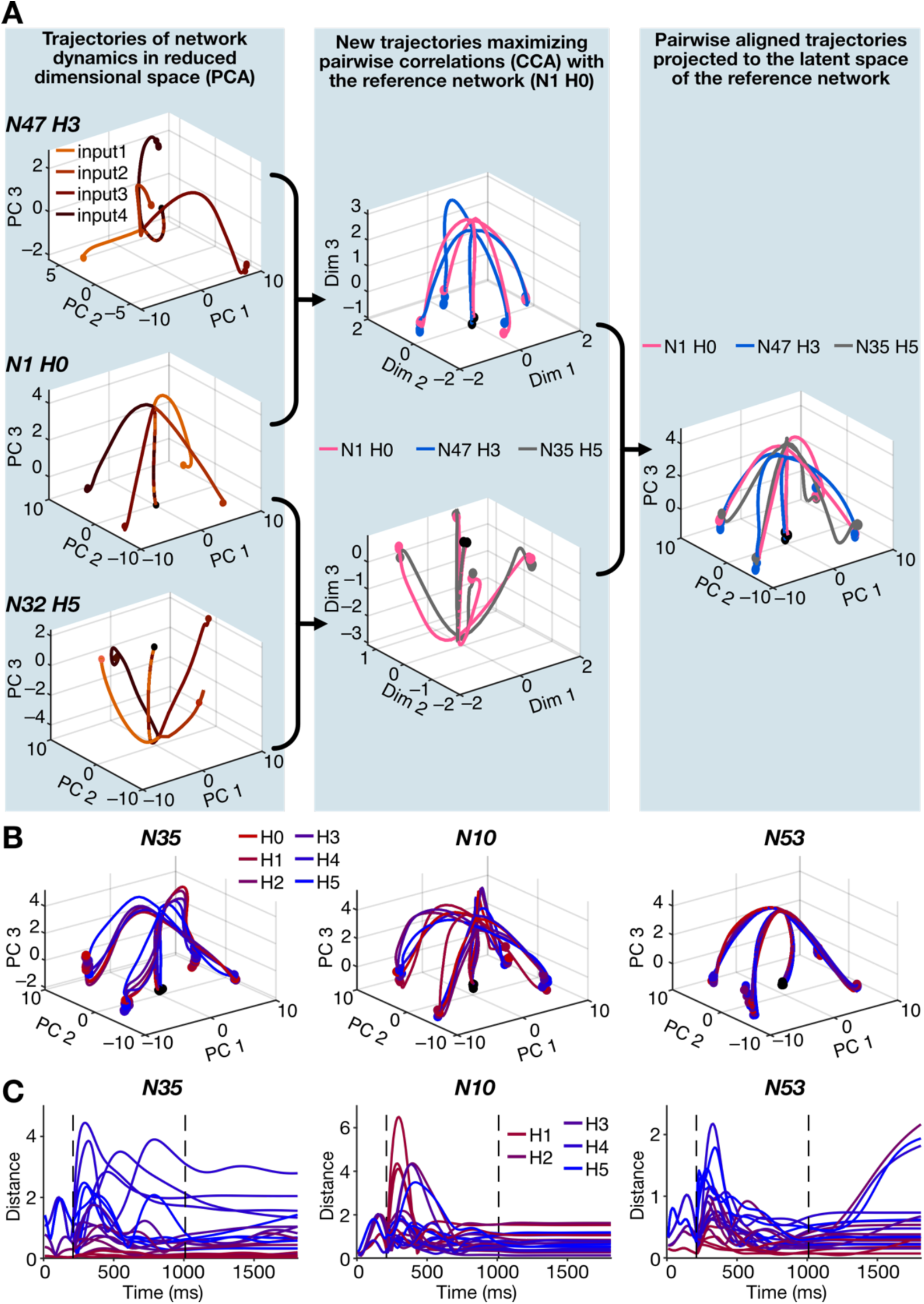
Pronounced network-to-network variability in latent dynamics of heterogeneous recurrent networks trained to perform a cognitive task. *A: Left,* A step-by-step representation of the process of aligning latent space dynamics of multiple networks. The dynamics of all 200 units in the network are represented by a trajectory in respective reduced dimensional spaces computed for each network using principal component analysis (PCA). *Center,* Using canonical correlation analysis (CCA), the latent space dynamics of each network and that of a reference network were transformed to a new coordinate space by maximizing the correlation between the trajectories in the new space in a pairwise manner. *Right,* The aligned trajectories in each CCA space were then projected back along the latent PCA space of the reference network bringing all the networks to a common reduced dimensional space, which enabled distance comparisons. *B:* The aligned latent space dynamics of three example networks, each shown for all 6 different levels of heterogeneity. *C:* Distance between trajectories, plotted as a function of time, of heterogeneous networks (H1–H5) with respect to the homogeneous network (H0) trajectory in the latent space.

When the same network with different levels of heterogeneities were compared, we found variability in task-related trajectories as well as the specific location of the input-specific fixed points in the aligned latent space (Fig. 3*B*). While some networks manifested pronounced variability in the dynamics across different levels of heterogeneities (*e.g.,* N35 and N10), there were networks (*e.g.,* N53) where the trajectories were less diverse across different levels of heterogeneities (Fig. 3*B*). We computed Euclidean distance between the latent-space trajectories of the heterogeneous H1–H5 networks and that of the homogeneous H0 network for each of the 53 network initializations. We plotted theses distances as functions of time for different networks and different with different levels of heterogeneities (Fig. 3*C*). These distance computations quantitively confirmed the visual observation that the variability in trajectories was higher in N35 and N10, while the distances between trajectories were comparatively lower for N53 (Fig. 3*C*). Importantly, in certain networks where the distances from the H0 trajectories increased with increasing level of heterogeneities (*e.g.*, N35), while in others the sequence reversed (*e.g.*, N10). In addition, the deviation in the fixed points (the final distance value at the end of the trial) depended on the level of heterogeneities and the type of input presented (*e.g.*, N35 and N53). Together, these trajectories provided examples for the pronounced network-to-network variability observed in the impact of training heterogeneities on network dynamics during task execution (Fig. 3*C*).

We assessed the distances of all heterogeneous networks (H1–H5) with their homogeneous counterpart (H0) for each of the different durations of the stimulus and response epochs across different input configurations (Fig. 4). Across different durations, we found network-to-network variability in latent space trajectories to initially increase and then decrease during the fixation epoch. We observed maximum variability in trajectories during the period that followed stimulus presentation (Fig. 4). Eventually, depending on how long the stimulus epoch was, variability in trajectories across heterogeneities reduced either during the response epoch (Fig. 4*A–B*) or during the stimulus epoch (Fig. 4*C–D*). Across durations, the variability was highest during a ∼300–400 ms duration, after which the trajectories converged to their respective fixed points with relatively minimal variability. As a population, the distance distributions across different durations of stimulus and response epochs were broadly comparable (Fig. 4). Therefore, for the rest of the analyses below, all distance-based calculations are shown for the longest duration trials.

**Figure 4.**
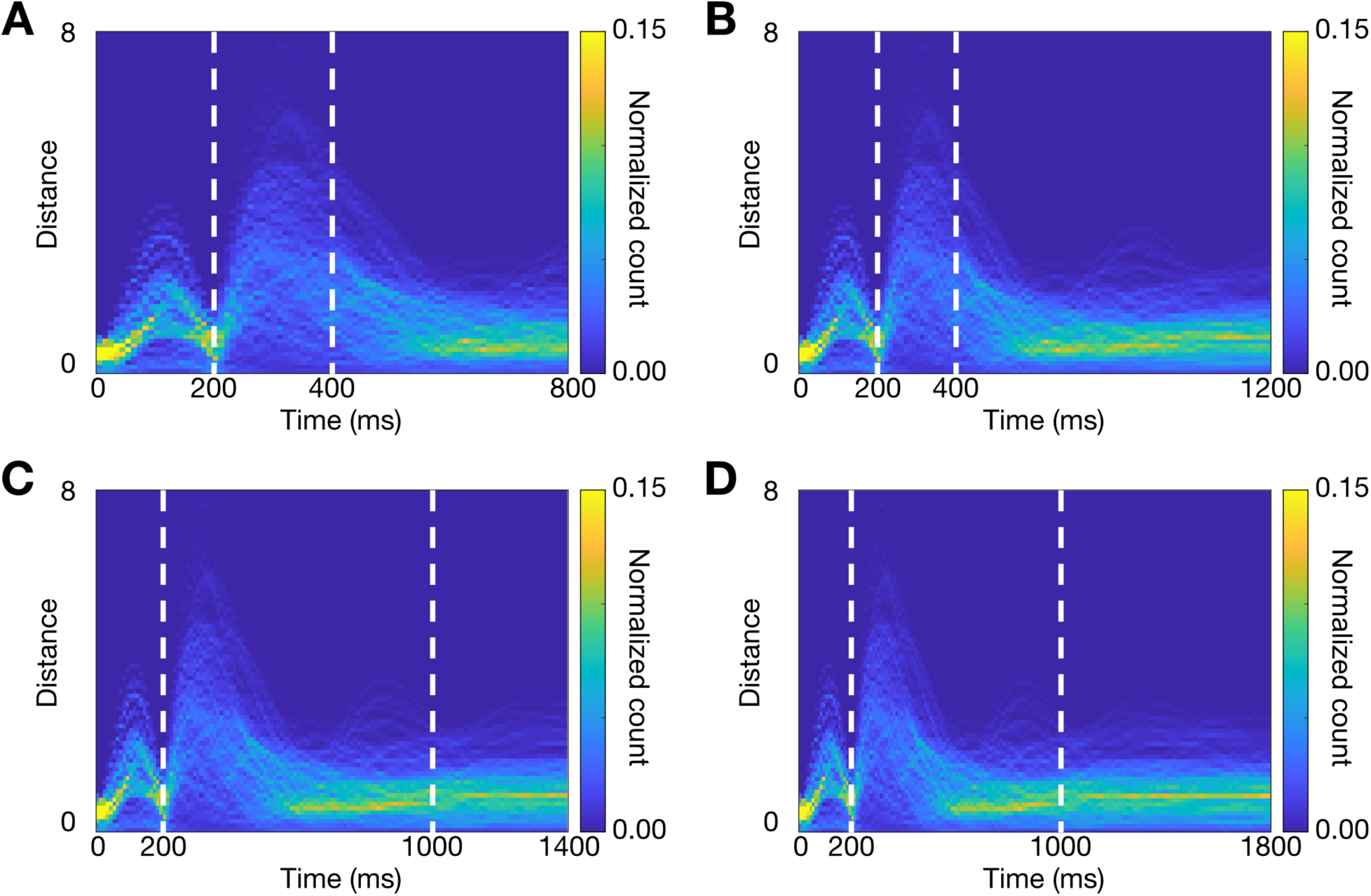
Network-to-network variability in latent dynamics of heterogeneous recurrent networks trained to perform a cognitive task across different epochs. *A–D:* Heatmaps of the distribution of Euclidean distances between all pairs of aligned latent space trajectories for all 318 networks. The fixation epoch remained constant at 200 ms for all panels. The stimulus epoch was 200 ms (A and B) or 800 ms (C and D). The response epoch was either 400 ms (A and C) or 800 ms (B and D). The variability in trajectories across networks especially during the stimulus period might be noted.

Together, these observations demonstrated that training heterogeneities in intrinsic properties impacted network dynamics in diverse ways even if they were trained with the same training algorithms, convergence criteria, and task specifications. The impact of training heterogeneities on network dynamics during task execution manifested pronounced variability in training performance (Fig. 2) as well as in task-execution dynamics (Figs. 3–4) across networks initialized with distinct sets of hyperparameters.

### Variable robustness of trained networks to post-training intrinsic heterogeneities

Biological neurons and networks could undergo changes post training, which could introduce heterogeneities in these properties. We tested the robustness of the trained networks to different such kinds of heterogeneities, specifically towards assessing the robustness of networks trained with different levels of training heterogeneities (H0–H5) to these post-training heterogeneities. We introduced post-training heterogeneities (referred to as P0–P5 to distinguish from training heterogeneities H0–H5) of different forms to each of the 318 networks at six different levels, starting at P0 representing no heterogeneity and P5 denoting the highest level of heterogeneity. The graded levels in different post-training heterogeneities were essential because (i) the exact level of perturbation in biological systems might be variable; and (ii) this approach allows for assessing the extent of different kinds of heterogeneities that the learnt system was robust to. For each of the several forms of post-training heterogeneities (Figs. 5–8), we performed all test trials and compared error levels and network dynamics across different levels of post-training heterogeneities for all 318 networks. The difference in errors as well as the distances measured in latent-space trajectories were computed for each of the P1–P5 heterogeneity levels with reference to the P0 network.

**Figure 5.**
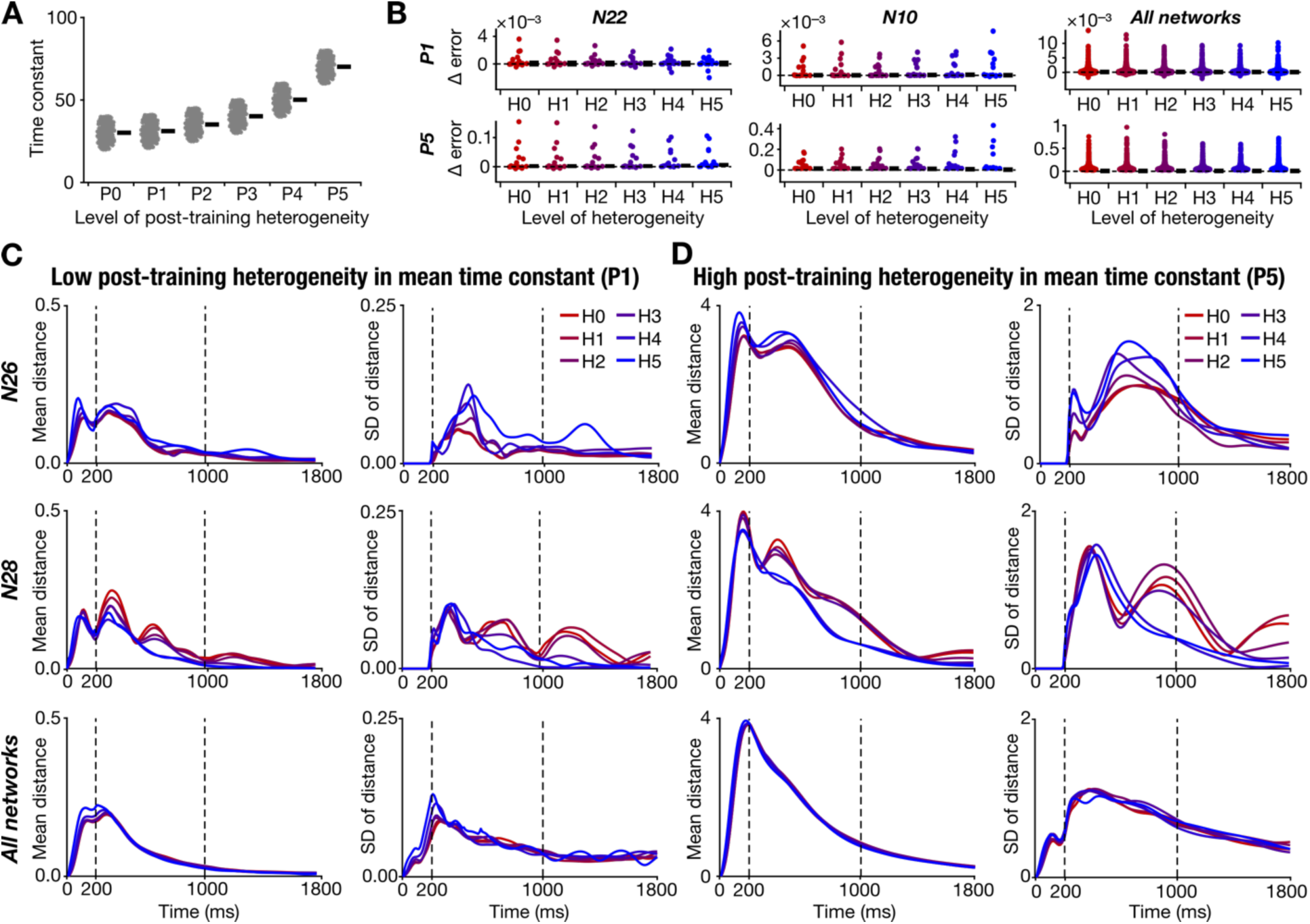
Pronounced network-to-network variability in the robustness of heterogeneous recurrent networks to post-training intrinsic heterogeneities. *A:* Distribution of time constants for six different levels of post-training heterogeneity (P0–P5). Note that the mean (black bar) increases as the level of post-training heterogeneities are increased, which is distinct from heterogeneities introduced during training (H0–H5; Fig. 1) where the mean was kept constant. *B:* Distribution of the difference in response errors (across different trials) with respect to P0 for low (P1; top row) and high (P5; bottom row) levels of post-training heterogeneity, introduced in networks trained with H0–H5 heterogeneities. The first column shows an example network (N22) that showed a reduction in error with increase in level of training heterogeneities (H0–H5). The second column depicts another network (N10) where the error increased in networks trained with higher level of heterogeneities. The third column summarizes the difference in errors across all 53 networks, spanning different trials. Black bars represent the median values. *C:* Mean (left column) and standard deviation (SD; right column) of the distance distribution (computed from trajectories in the latent space) with respect to P0 for low level of post-training heterogeneity (P1). Each panel shows plots for networks trained with different levels of heterogeneity (H0–H5). The first row shows an example network (N26) where the mean distance and the associated variability increased with increasing level of training heterogeneities. The second row depicts another network (N28) where the mean distance and the associated variability decreased with increasing level of training heterogeneities. The third row depicts mean distance and the associated variability across all 53 networks. *D:* Same as *C* but for P5 instead of P1.

We studied the impact of post-training heterogeneities in intrinsic properties of recurrent units after training was complete. Specifically, we tested two types of post-training intrinsic heterogeneities in time constant distributions. First, the extent of heterogeneities in terms of the distance of time constants from the mean was kept constant, while the mean time constant was gradually shifted (Fig. 5*A*). Across networks, we observed an increase in error in trial performances (Fig. 5*B*) as well as increase in distances of latent-space trajectories (Fig. 5*C* vs. Fig. 5*D*), both computed with reference to the P0 trials, with increase in post-training heterogeneities from P1 to P5. With increase in level of post-training heterogeneities from P1 (Fig. 5*C*) to P5 (Fig. 5*D*), we found an increase in both mean distance and the variability (SD) across different types of inputs.

We then asked if the robustness of the network in terms of error performance or network dynamics to post-training heterogeneities (P0–P5) was dependent on the level of intrinsic heterogeneities used during training (H0–H5). We found that in certain networks (*e.g.*, N22), the change in error (from P0) decreased with increase in level of training heterogeneities (from H0 to H5), whereas in other networks (*e.g.*, N10) the change in error increased from H0–H5. Similarly, there were certain networks (*e.g.*, N28) where the mean and variance in trajectory distances decreased with increasing training heterogeneities, whereas in other networks (*e.g.*, N26), the opposite was true. As a consequence of pronounced variability within networks (across trials) and across networks, the overall differences in errors (Fig. 5*B*; All networks) as well as the mean and SD in distances (Fig. 5*C–D*; All networks) were comparable across H0–H5 levels of training heterogeneities.

In a second scenario (Supplementary Fig. S4*A*), we retained the mean to be the same (30 ms) with the variance of post-training heterogeneities changing across P0–P5 (with P0 representing zero variance). As this was identical to how H0–H5 were defined (Fig. 1), a network trained with Hx showed minimal error for Px (Supplementary Fig. S4*B*). Therefore, in this case, errors and distances were computed with reference to the heterogeneity level that the specific network was trained with, rather than with the respective P0. The changes in error were minimal for the heterogeneity level that the network was trained with and increased with increasing distance from the level that it was trained with (Supplementary Fig. S4*B*). A similar observation held for the mean and SD of distances in the latent-space distance trajectories (Supplementary Fig. S4*C– D*). Importantly, there was network-to-network variability in terms of error performance in H0– H5 networks (Supplementary Fig. S4*B*) as well as in the mean and variance of latent-space distance trajectories (Supplementary Fig. S4*C–D*).

### Variable robustness of trained networks to post-training synaptic heterogeneities

Learnt synaptic weights in a biological network could change because of training for additional tasks or homeostatic mechanisms or non-specific perturbations. To assess the impact of such post-training synaptic changes, we added Gaussian jitter with progressively increasing variance (P0– P5) to the trained synaptic weights (Fig. 6*A*) and evaluated network performance and dynamics of all 318 networks. Performance errors were high for most networks and got progressively worse with increasing levels of synaptic jitter (Fig. 6*B*). There were few networks where the difference in errors with synaptic jitter showed an increase (*e.g.*, N11) or decrease (*e.g.*, N8) from H0 to H5 networks (Fig. 6*B*). On the average, error distribution across H0–H5 was comparable for P1 and P5 (Fig. 6*B*). Importantly, the mean and the variability in the distances between trajectories of P1– P5 networks, compared to their P0 counterparts, were high even during the response epoch (Fig. 6*C–D*). Deviations from the P0 network dynamics progressively increased from P1 to P5 (Fig. 6*C* vs. Fig. 6*D*). These observations indicated that the networks were unable to converge to the required fixed points when post-training synaptic jitter was introduced (Fig. 6*C–D*). Network-to-network variability in the robustness of network dynamics on pre-training heterogeneities was observed in a few networks, with some showing increased distances (N22) while others showed a reduction (N16) through H0–H5 (Fig. 6*C–D*).

**Figure 6.**
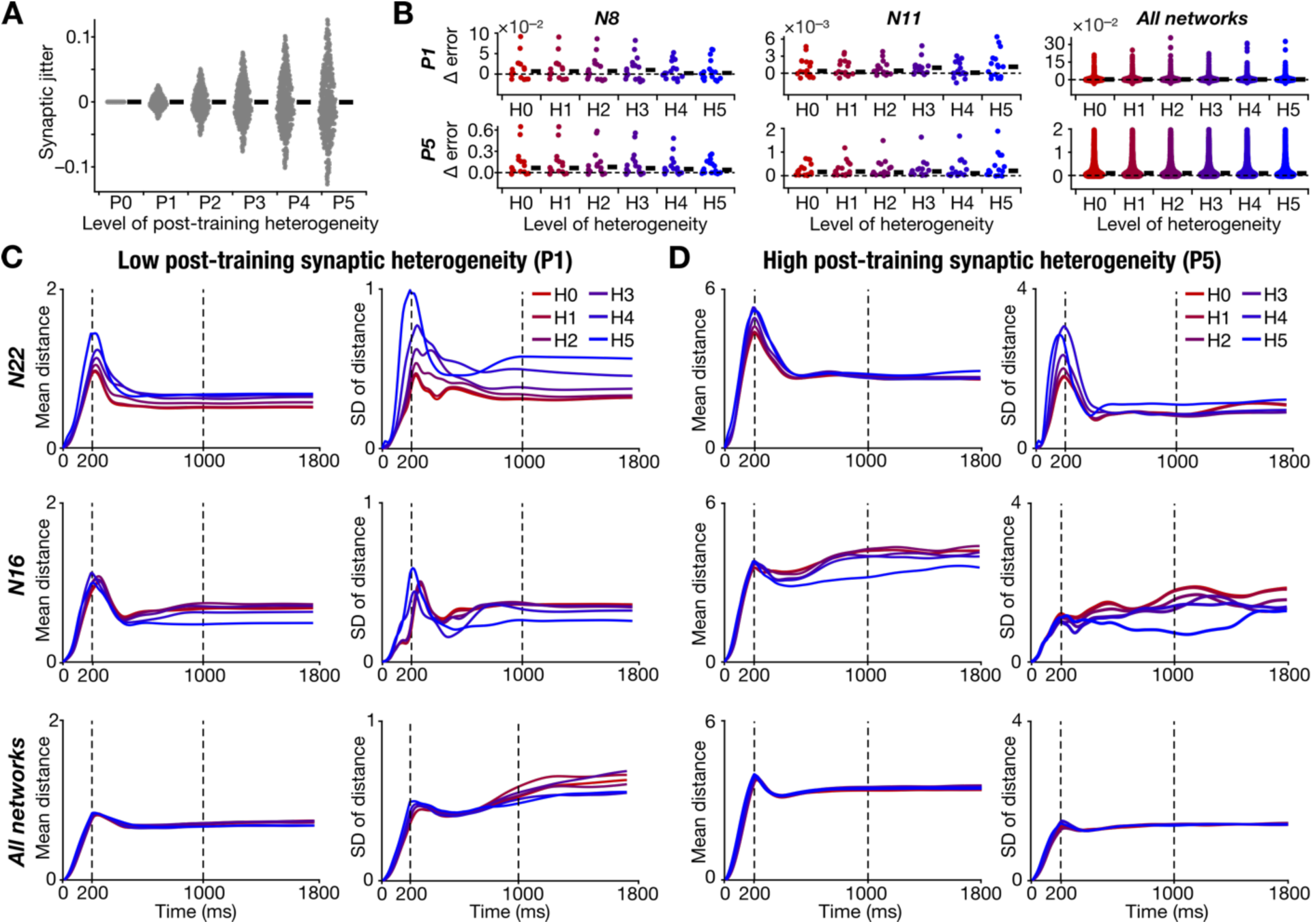
Pronounced network-to-network variability in the robustness of heterogeneous recurrent networks to post-training synaptic jitter. *A:* Distribution of synaptic jitter (deviation from trained values) for six different levels of post-training heterogeneity (P0–P5). *B:* Distribution of the difference in response errors (across different trials) with respect to P0 for low (P1; top row) and high (P5; bottom row) levels of post-training heterogeneity, introduced in networks trained with H0–H5 heterogeneities. The first column shows an example network (N8) that showed a reduction in error with increase in level of training heterogeneities (H0–H5). The second column depicts another network (N11) where the error was high in networks trained with higher level of heterogeneities. The third column summarizes the difference in errors across all 53 networks, spanning different trials. Black bars represent the median values. *C:* Mean (left column) and standard deviation (SD; right column) of the distance distribution (computed from trajectories in the latent space) with respect to P0 for low level of post-training heterogeneity (P1). Each panel shows plots for networks trained with different levels of training heterogeneity (H0–H5). The first row shows an example network (N22) where the mean distance and the associated variability increased with increasing level of training heterogeneities. The second row depicts another network (N16) where the mean distance and the associated variability decreased with higher level of heterogeneities introduced during training. The third row depicts mean distance and the associated variability across all 53 networks. *D:* Same as *C* but for P5 instead of P1. Note that in panels C and D, the mean distance was high even during the response window, indicating the deleterious impact of synaptic jitter on network performance.

### Variable robustness of trained networks to post-training heterogeneities in the initial activity of the network and activity perturbations

We tested the robustness of the network to two types of post-training heterogeneities in initial activity patterns at the beginning of the trials (𝒙(0)). First, the extent of heterogeneities in terms of the initial activity patterns from the mean was kept constant while gradually shifting the mean activity value from P0–P5 (Fig. 7*A*). Changes in the initial state affected few trials in certain networks, but mostly the changes in error values were minimal across networks across different levels of post-training heterogeneities (Fig 7*B*). Network-to-network variability in the dependence of performance error on training heterogeneities was prevalent (Fig. 7*B*), with some networks showing less error for H0 compared to H5 (*e.g.*, N26) and others showing the opposite trend (*e.g.*, N41). The network dynamics manifested deviations in network trajectories, which progressively increased through P1–P5, with changes in the initial state of the network. However, these deviations were predominantly limited to the first half of the trial period (Fig. 7*C–D*). In most cases, the network converged to the same final fixed points, thereby minimizing the final errors during the response epochs (Fig. 7*C–D*). We did observe some networks showing more deviation in network dynamics for high levels of training heterogeneities (*e.g.*, N40) whereas others showed enhanced deviations for low levels of training heterogeneities (*e.g.*, N25) for both P1 (Fig. 7*C*) and P5 (Fig. 7*D*) levels of post-training heterogeneities. In a second scenario (Supplementary Fig. S5), we kept the mean activity to be the same with the range of post-training heterogeneities changing across P0–P5. Our observations here were very similar to the scenario where we changed the mean activity, whereby the deviations in dynamics (which did show network-to-network variability) had negligible impact on the response epoch and final outcomes (Supplementary Fig. S5).

**Figure 7.**
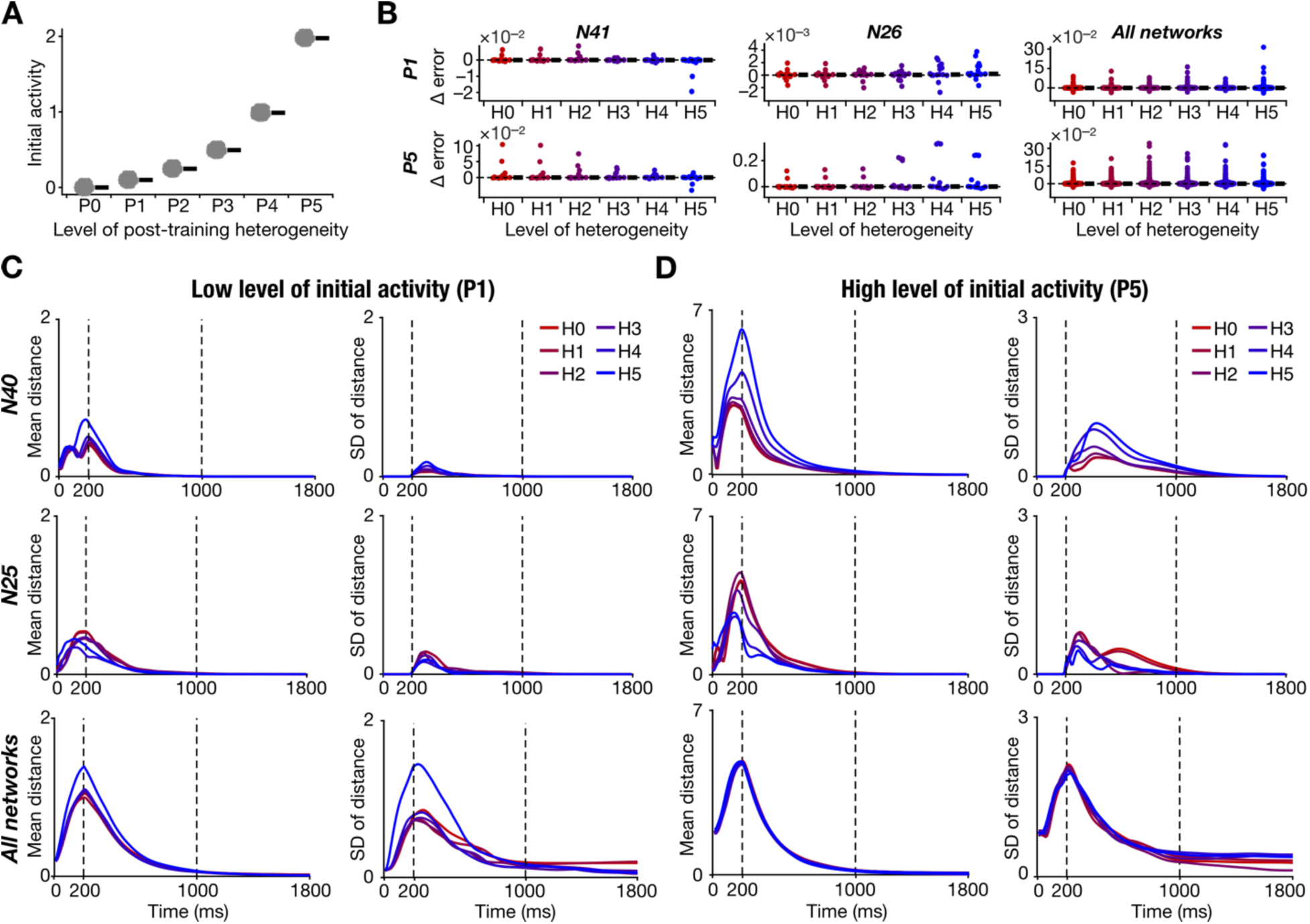
Pronounced network-to-network variability in the robustness of heterogeneous recurrent networks to post-training shift in network activity at task initiation. *A:* Distribution of network activity range in all recurrent units at network initiation for six different levels of post-training heterogeneity (P0–P5). Note the shift in the mean value of the initial activity distribution with increasing in level of post-training heterogeneity. *B:* Distribution of the difference in response errors (across different trials) with respect to P0 for low (P1; top row) and high (P5; bottom row) levels of post-training heterogeneity, introduced in networks trained with H0–H5 heterogeneities. The first column shows an example network (N41) that showed a reduction in error with increase in level of training heterogeneities (H0–H5). The second column depicts another network (N26) where the error was high in networks trained with higher level of heterogeneities. The third column summarizes the difference in errors across all 53 networks, spanning different trials. Black bars represent the median values. *C:* Mean (left column) and standard deviation (SD; right column) of the distance distribution (computed from trajectories in the latent space) with respect to P0 for low level of post-training heterogeneity (P1). Each panel shows plots for networks trained with different levels of training heterogeneity (H0–H5). The first row shows an example network (N40) where the mean distance and the associated variability increased with increasing level of training heterogeneities. The second row depicts another network (N25) where the mean distance and the associated variability decreased with higher level of heterogeneities introduced during training. The third row depicts mean distance and the associated variability across all 53 networks. *D:* Same as *C* but for P5 instead of P1.

We tested the network by introducing activity perturbations, Δ (which were used only during training trials, but not during test trials), at frequencies in between 0 and 10 Hz defining P0–P5 (Supplementary Fig. S6*A*). Since the networks were trained with low frequency Δ, we expected that the network would be robust to low-frequency activity perturbations. We found the difference in errors to be low (<0.4) across all levels of training and post-training heterogeneity (Supplementary Fig. S6*B*). Network-to-network variability in the dependence on the level of training heterogeneities was observed, with some networks showing increasing error through H0– H5 (*e.g.*, N13) while others showed a reduction (*e.g.*, N3). The distance between network trajectories was high due to random activity perturbations (Supplementary Fig. S6*C–D*). On the average across networks, the deviation in network dynamics peaked at the start of stimulus epoch and the maximal deviations were not markedly different across H0–H5 levels of training heterogeneity (Supplementary Fig. S6*C–D*). However, there were networks like N20 where the deviations and their variance were higher for H5 than for H0, or like N53 where the deviations and their variance were higher for the H0 network than others (Supplementary Fig. S6*C–D*).

Together, we found large deviations in network dynamics with changes in network initialization, coupled with pronounced network-to-network variability in the dependence on training heterogeneities. However, these deviations did not affect the final responses as the network converged to the same fixed points during the response epoch of most trials across P1–P5 levels of post-training heterogeneities in network initialization (Fig. 7, Supplementary Fig. S5). With activity perturbations, we found pronounced network-to-network variability in robustness of network performance and its dependence on training heterogeneities, although overall errors in performance were low across different frequencies of perturbation (Supplementary Fig. S6).

### Variable robustness of trained networks to changes in stimulus epoch duration

During the training process, we had used a stimulation epoch duration of either 200 ms or 800 ms (Fig. 1). We tested the robustness of the learnt networks to variations in the stimulus epoch duration from 800 ms to 50 ms spanning P0–P5 levels of post-training heterogeneities (Fig. 8*A*). The changes in errors were not high for H0 to H5 networks when epoch durations remained higher than 100 ms (Fig. 8*B*). However, for epoch durations less than 100 ms, the errors were high irrespective of the level of training heterogeneity (Fig. 8*B*). Given that the design of these trials involved variable stimulus epoch durations (through P0–P5), computation of distances between the network trajectories (comparing P1–P5 with P0 dynamics) was feasible only for the response epoch (Fig. 8*C–D*). The distance values at the beginning of the response epoch progressively increased with reduction in stimulus epoch duration (Fig. 8*C* vs. Fig. 8*D*). However, across P1–P5, the deviation in network trajectories sharply declined and showed convergence in many networks, although the variability was high with shorter stimulus epoch durations (Fig. 8*C–D*). Network-to-network variability was prevalent in terms of how robustness to stimulus epoch duration depended on H0– H5 levels of training. Whereas some networks (*e.g.*, N23) showed lower distances for H5 *vs*. H0, others (*e.g.*, N52) showed the opposite dependence. Nevertheless, the average distance and the associated variance across all networks did not vary much across H0 to H5 (Fig. 8*C–D*). Together, decreasing the stimulus epoch reduced the accuracy of network performance and enhanced deviations in network dynamics and associated variability. There was pronounced network-to-network variability in how the robustness to changes in stimulus duration depended on training heterogeneities.

**Figure 8.**
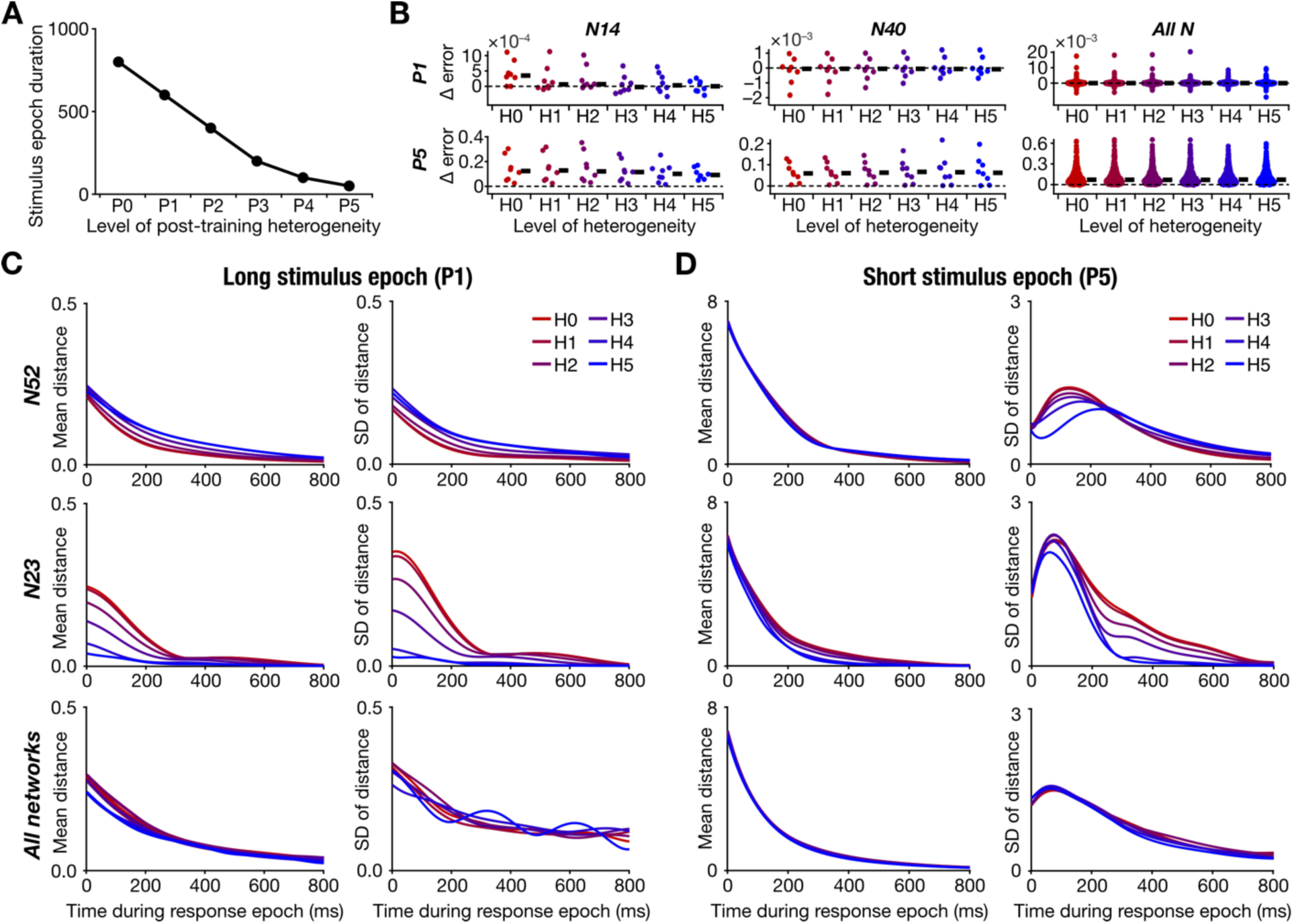
Pronounced network-to-network variability in the robustness of heterogeneous recurrent networks to post-training shift in the stimulus epoch duration. *A:* Value of stimulus epoch duration for six different levels of post-training heterogeneity (P0–P5). *B:* Distribution of the difference in response errors (across different trials) with respect to P0 for low (P1; top row) and high (P5; bottom row) levels of post-training heterogeneity, introduced in networks trained with H0–H5 heterogeneities. The first column shows an example network (N14) that showed a reduction in error with increase in level of training heterogeneities (H0–H5). The second column depicts another network (N40) where the error was high in networks trained with higher level of heterogeneities. The third column summarizes the difference in errors across all 53 networks, spanning different trials. Black bars represent the median values. *C:* Mean (left column) and standard deviation (SD; right column) of the distance distribution (computed from trajectories in the latent space) with respect to P0 for low level of post-training heterogeneity (P1). Each panel shows plots for networks trained with different levels of training heterogeneity (H0–H5). The first row shows an example network (N52) where the mean distance and the associated variability increased with increasing level of training heterogeneities. The second row depicts another network (N23) where the mean distance and the associated variability decreased with higher level of heterogeneities introduced during training. The third row depicts mean distance and the associated variability across all 53 networks. *D:* Same as *C* but for P5 instead of P1. For panels C and D, the mean distance and the associated variability are depicted for the response epoch, as distance computation during the stimulus epoch was impossible, given the differences in stimulus epoch duration across the different post-training heterogeneity levels.

## DISCUSSION

We designed and employed a systematic framework for quantitative assessment of the impact of neural heterogeneities on recurrent networks trained to perform a cognitive task. We introduced six different levels of intrinsic heterogeneities during training in a population of 53 networks, each obtained by random sampling of network hyperparameters. We assessed each of these 318 trained networks in terms of their training performance, response dynamics, and robustness to six different levels of several post-training heterogeneities. We demonstrated that intrinsic heterogeneities impacted network dynamics in diverse ways even if they were trained with the same training algorithm, convergence criteria, and task specifications. Our analyses unveiled pronounced network-to-network variability in terms of the number of training trials required for learning, the error values associated with specific trials, and the dependencies of trial numbers and the error values on the level of heterogeneity (Fig. 2) as well as on network parameters (Supplementary Fig. S1). Furthermore, the impact of training heterogeneities on network dynamics during task execution also manifested pronounced variability across networks (Figs. 3–4). Our analyses also revealed a prominent impact of post-training heterogeneities on performance errors and network dynamics, with progressive increases in errors as well as in trajectory deviations with graded increases in post-training heterogeneities. Importantly, our analyses demonstrated pronounced variability in the robustness, to different levels of post-training heterogeneities, of networks trained with different levels of training heterogeneities. Specifically, whereas certain networks showed enhanced robustness to post-training heterogeneities when training heterogeneities were low, others showed better robustness when training heterogeneities were high (Figs. 5–8, Supplementary Figs. S4–S6). Of all the post-training heterogeneities tested, we found synaptic jitter (Fig. 6) to be the most detrimental post-training heterogeneity, which severely affected all networks across all training heterogeneity levels even during the response epochs.

### The need for a population of networks approach in studying the impact of neural-circuit heterogeneities

The pronounced network-to-network variability in the impact of heterogeneities in training performance and in task-execution dynamics critically emphasize the need for using a population of networks approach to study neural-circuit heterogeneities. The use of a single network would have biased conclusions, showing that heterogeneities had either a beneficial or a deleterious impact on performance. The use of a population of networks allowed us to identify different networks that manifested diverse dependencies on heterogeneities, in training performance and task dynamics. The importance of using a population of networks approach has been strongly emphasized by several studies to account for variability as well as the role of distinct components in implementing network function (Prinz et al., 2004; Schaeffer et al., 2024; Calabrese and Marder, 2025). Our analyses strongly emphasize the need for the use of population of networks approach in studying the impact of heterogeneities on recurrent networks implementing cognitive tasks.

An important aspect of the population of networks approach is the impact of perturbations on network performance. It has been repeatedly observed that networks performing the same tasks, but with variable sets of components respond differently to the same perturbations (Mishra and Narayanan, 2021a; Ratliff et al., 2021; Gorur-Shandilya et al., 2022; Marder et al., 2022; Alonso et al., 2023; Marom and Marder, 2023; Schapiro and Marder, 2024; Calabrese and Marder, 2025). The variable impact of these perturbations has also been used to assess the dynamics of dysfunction in these neural circuits. In this context, our analyses demonstrate pronounced network-to-network variability in the impact of different forms of post-training heterogeneities on network performance. Whereas certain networks manifested strong dependence on training heterogeneities, others showed negligible impact on the level of training heterogeneities. The diversity in the dependence of resilience to different forms of post-training heterogeneities could be employed as a handle on how different networks accomplish the task and what contributions training heterogeneities make to this resilience. Together, our framework strongly recommends the training and assessment of a population of networks with different hyperparameters in studying the impact of different forms of neural-circuit heterogeneities on task performance, dynamics, and resilience to post-training heterogeneities.

### Diversity in the impact of heterogeneities across networks and their functional outputs

The study of neural heterogeneities has revealed a diversity of roles played by heterogeneities in affecting circuit function. Heterogeneities have been shown to be beneficial in terms of performance and resilience of decorrelating networks (Padmanabhan and Urban, 2010; Tripathy et al., 2013; Mishra and Narayanan, 2019, 2021a) as well as in recurrent networks that have been trained to perform certain classification tasks (Perez-Nieves et al., 2021). In contrast, heterogeneities have been shown to hamper performance of continuous attractor networks (CAN), although the addition of a negative feedback loop to individual units stabilize heterogeneous CAN models (Mittal and Narayanan, 2021; Gast et al., 2024). Our analyses here extend this diversity of impact by demonstrating pronounced network-to-network variability in the impact of heterogeneities on training, task-execution dynamics, and resilience of recurrent networks performing a cognitive task. Our framework involved the training of recurrent networks with different hyperparameters and with different levels of training heterogeneities. Our assessment of trained networks spanned training performance, task execution, and resilience to several levels of different post-training heterogeneities with a variety of performance metrics. In conjunction with other studies mentioned above, our analyses suggest that the diversity in the impact of heterogeneities is critically dependent on several factors. Factors that govern such diversity include the kind of network and its architecture, the type of model used for individual units, network hyperparameters, the specific task that is performed by the network and its complexity, the type of heterogeneity being introduced, the degree of heterogeneity introduced, the specific methodology used to introduce heterogeneity, the metrics used to evaluate performance. In addition to these our analyses also emphasize that the level and form of post-training heterogeneities also need to be explicitly accounted for in assessing resilience of heterogeneous networks to perturbations. Overall, our analyses emphasize the critical need to account for each of these factors in assessing the impact of neural-circuit heterogeneities on circuit function. Without accounting for these factors, conclusions are bound be biased by the choices made with reference to each of these different factors that govern the diversity in the impact of heterogeneities on circuit function.

### A complex system interpretation of heterogeneities in neural circuits

How do we reconcile this diversity and variability in the dependence of network function on component heterogeneities? Our analyses suggest the complex systems framework as an ideal substrate for analyzing the impact of heterogeneities on neural circuits and their performance. Complex systems are systems where several functionally specialized subsystems interact to yield collective functional outcomes (Watts and Strogatz, 1998; Edelman and Gally, 2001; Milo et al., 2002; Kim and Wilhelm, 2008). With specific reference to biological neural circuits and the different forms of heterogeneities that manifest there, each neuron could be considered as a functionally specialized subsystem and the synaptic connectivity defining specific interactions (Luo, 2021). Complex systems are defined by two key attributes. First, the interactions among subsystems of a complex system are neither fully determined nor completely random. This intermediate level of randomness is characterized by network motifs, defined as subnetworks that appear more frequently than expected in random networks. With reference to neural circuits and their heterogeneities, arbitrarily random synaptic connectivity with random neuronal heterogeneities wouldn’t accomplish a given task. This may be readily inferred from the high pre-training errors in task execution (Fig. 2). Thus, learning could be considered as a mechanism that identifies these motifs (the non-random interactions) that are essential for execution of the specific task in hand (Driscoll et al., 2024; Kuan et al., 2024).

The second key feature of complex systems, which rules out the “completely determined” extreme in complex systems, is degeneracy, defined as the ability of multiple combinations of distinct subsystems to achieve the same collective function (Edelman and Gally, 2001; Prinz et al., 2004; Rathour and Narayanan, 2019; Seenivasan and Narayanan, 2022; Albantakis et al., 2024). In a heterogeneous neural circuit, each neuron is endowed with disparate intrinsic properties and neurons are connected by weight distributions that are non-uniform. Thus, the ability of disparate sets of functionally specialized subsystems interacting with each other in heterogeneous ways to yield the same collective function (implement the task) constitutes a form of degeneracy. The presence of several such combinations, elucidated by different network initializations with disparate forms of intrinsic and synaptic heterogeneities, that are successfully able to learn the task (Fig. 2) with disparate network dynamics (Fig. 3–4) fulfils this.

Therefore, *individual* neural heterogeneities shouldn’t be seen through the prism of whether they are beneficial or detrimental to circuit function. Instead, they should be interpreted from the perspective how specific *combinations* of disparate forms of heterogeneities come together to implement a specific function. When each of the different components of the network are assessed individually, they manifest widespread heterogeneities. But, when specific forms of heterogeneities interact with each other, they are able to accomplish the collective function of the complex system that they define. For example, in our analysis, disparate combinations of interactions among specific synaptic weights with specific intrinsic properties (each of which is heterogeneous by itself within that network) implements collective function. Even in scenarios where heterogeneities have been shown to be deleterious, the presence of other components with appropriate heterogeneities in them have negated the deleterious impact of heterogeneities. For instance, different forms of heterogeneities have been shown to negatively impact continuous attractor networks (Mittal and Narayanan, 2021; Gast et al., 2024). However, the incorporation of an additional heterogeneous component that implements a slow negative feedback loop, either through inhibition or through ionic currents in individual neurons, could stabilize these networks towards accomplishing their collective function (Mittal and Narayanan, 2021; Gast et al., 2024). Therefore, within the complex systems framework, we argue that the focus shouldn’t be on heterogeneities in individual components. Instead, the emphasis must be on the global structure of heterogeneities across all components and how they interact with each other towards achieving collective function (Goldman et al., 2001; Prinz et al., 2004; Rathour and Narayanan, 2019; Mishra and Narayanan, 2021b; Seenivasan and Narayanan, 2022; Mittal and Narayanan, 2024).

### Limitations and future directions

Our analyses here was to develop a framework for assessing the impact of neural heterogeneities on circuit function, with networks endowed with a simple form of heterogeneity trained with one of the simplest of cognitive tasks. These analyses should be extended to networks trained to perform multiple tasks, each with more complexity than the task that we had employed here. Such analyses also could bring in other biological attributes such as networks with separate excitatory and inhibitory neuronal populations, single units with more complexity reflecting biological neurons with active dendrites, concomitant plasticity of intrinsic and synaptic properties, and connectivity (local as well as afferent) that reflects specific biological networks (Song et al., 2016; Mishra and Narayanan, 2019; Luo, 2021; Mishra and Narayanan, 2021a, b; Perez-Nieves et al., 2021; Khona and Fiete, 2022; Chavlis and Poirazi, 2024; Pagkalos et al., 2024). Such incorporations would increase the number of places where heterogeneities could be introduced at different levels, providing avenues for assessment of training performance, task-execution dynamics, and resilience to perturbation when networks are trained with these heterogeneities. An important avenue for further exploration on the impact of heterogeneities is in continual learning systems, especially where the compositionality of dynamic motifs aids continual learning of different tasks in the same network (Yang et al., 2019; Driscoll et al., 2024). An important line of future research would be to assess the quantitative impact of different neurological disorders on neural-circuit heterogeneities and how such changes in the level or form of heterogeneities affect circuit function. Although most analyses focus on the average shifts in specific quantities, given the diverse and strong impact of neural-circuit heterogeneities on circuit function, it is essential to quantify this across different measurements and ask if altered heterogeneities contribute to circuit function (Rich et al., 2022).

## METHODS

### Network architecture

A fully connected recurrent network model of 𝛮 (=200) units (Fig. 1) was used for all analyses. The dynamical equation that governed network activity was (Miconi, 2017):

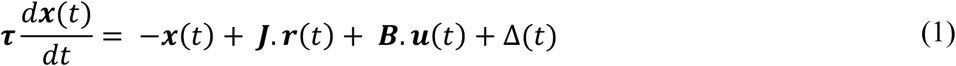

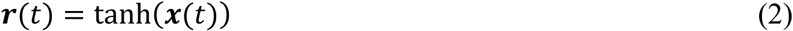

where 𝒙(𝑡) represented the *N*-dimensional activity vector spanning all units, 𝒓(𝑡) defined the response vector of the network units, and 𝒖(𝑡) represented the *M*-dimensional feed-forward input vector at time 𝑡. 𝑱 depicted an 𝛮 × 𝛮 matrix of recurrent connection weights, initialized from a normal distribution of mean 0 and variance 𝑔^2^/𝑁. 𝑩 is an 𝛮 × 𝑀 matrix of input connection weights initialized from a uniform distribution in the range [−0.01, 0.01]. 𝝉 denoted an *N*-dimensional vector of time constants assigned to each unit in the network. To introduce trial to trial variability during training, random exploratory perturbations Δ, were independently introduced to the activity of each unit at an average frequency of 3 Hz. The amplitude of these perturbations were sampled from a uniform distribution that spanned [–0.5,0.5] and occurred randomly. One of the 𝛮 units was designated as the output unit of the network whose response is evaluated as the network’s response.

The vector 𝒖(𝑡) was made of three different types of inputs: 𝒖 = (𝑢*_fix_*, 𝒖*_stim_*, 𝒖*_task_*)E. The fixation input 𝑢*_fix_* distinguishes the fixation period of the cognitive task from the response period. At the beginning of a trial, 𝑢*_fix_* was assigned a value of 1, indicative of the fixation period. Switching the value of 𝑢*_fix_* to 0 signals the beginning of the response epoch, during which the network response was evaluated. Stimulus input vector 𝒖*_stim_* = (𝑢*_stim_*_1_, 𝑢*_stim_*_2_) is 2-dimensional and carried task-specific stimulus information to all units of the network. Each input 𝑢*_stim1_* and 𝑢*_stim2_* switched between one of three values: –1, 0, or 1, with 0 indicating the absence of any stimulus and –1/1 representing one of two distinct kinds of inputs that the network was presented with. 𝒖*_task_*(𝑡) is a 6-dimensional one-hot vector with each dimension specifying a different task. For any trial, only the task input corresponding to the current task was set at 1, while the others remained at 0. For all experiments reported in this study, the dimension for the “Go task” was set to 1 and the rest were set to zero. In our simulations, a first-order Euler approximation in time steps of Δ𝑡 for each unit 𝑖 was used for implementing the dynamical system represented in equation (1):

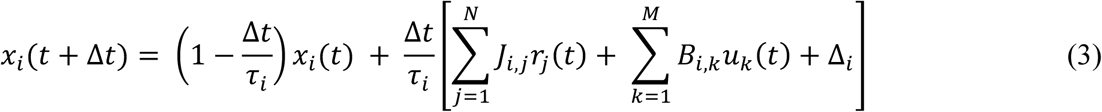

where 𝛮 = 200, 𝑀 = 9, 𝑔 = 1.5 and Δ𝑡 = 1 ms. The initial activity pattern 𝒙(0) was set by sampling a uniform distribution spanning [–0.1,0.1]. To ensure that the units received information about the stimulus inputs (𝒖*_stim_*), the column vectors in 𝑩 corresponding to the stimulus inputs were scaled up by a factor of 100. Four arbitrarily chosen units were maintained at a constant activation of 1 to act as bias inputs to other units in the network.

### Incorporating training-heterogeneity in the model

The time constant associated with individual units was initialized as a vector to incorporate intrinsic heterogeneity into the network. 𝝉 was initialized from a uniform distribution in range [𝜏_min_, 𝜏_max_]. Consistent with our earlier analyses on heterogeneities (Mittal and Narayanan, 2021, 2022), we introduced heterogeneities in a graded fashion rather than using a binary homogeneous *vs.* heterogeneous distinction in networks. In implementing this gradation, we introduced six levels of heterogeneities in unit time constants. For the six levels of heterogeneities H0–H5 in that order, the ranges of time constants [𝜏_min_, 𝜏_max_] were defined by 𝜏_min_ = 30, 29, 25, 20, 10, 1 and 𝜏_max_ = 30, 31, 35, 40, 50, 59 (Fig. 1). The homogeneous level (H0) corresponded to a network where all units were assigned an identical time constant value of 30 ms. The level of heterogeneities progressively increase through H1 to H5 as indicated by the respective values of 𝜏_min_and 𝜏_max_, translating to a progressive increase in the range of time constants always centered at 30 ms (Fig. 1). The highest heterogeneity level (H5) was defined by units being randomly assigned to time constant values in the range 1–59 ms. The choice of the range 1–59 ms was to ensure that the time constant doesn’t become zero for any unit and that the mean time constant across all heterogeneities was fixed at 30 ms.

To avoid biases introduced by the use of a single network for all analyses, we used a population of networks approach throughout our analyses to assess potential network-to-network variability in training performance and response dynamics. Specifically, the major hyperparameters of our network are: the seed value that determines the initial values of the connectivity matrices 𝑱 and 𝑩, the seed value that determines the initial values of the activity vector 𝒙(0), the seed value that determines the values of time constants from their respective distributions (in heterogeneous networks), and the seed value governing the pattern of exploratory perturbation Δ. Instead of using one set of initial conditions associated with these hyperparameters, we created 53 distinct networks (N1–N53) by picking 53 different values for these hyperparameters. Each of these 53 distinct networks were initialized with the 6 different intrinsic heterogeneities (H0–H5), together providing a total of 53 × 6 = 318 total networks that were trained for the cognitive task. The presence of a population of networks, each trained with different levels of heterogeneities, allowed us to quantitatively assess network-to-network variability in the impact of heterogeneities on training performance and response dynamics.

### Cognitive task description

Each of the 318 networks were trained on a memoryless Go task (Fig. 1). Each trial of the task consisted of three epochs. The *fixation epoch* was 200 ms long, during which 𝑢*_fix_* = 1. At the beginning of the *stimulation epoch*, a single stimulus was presented through either 𝑢*_stim_*_1_ or 𝑢*_stim_*_2_inputs. Specifically, one of 𝑢*_stim_*_1_or 𝑢*_stim_*_2_ was randomly set to 1 or –1 and remained at that stimulus value for the rest of the entire trial duration. The stimulation epoch duration switched randomly between 200 ms and 800 ms to ensure that the network learned the task and not the specifically timed responses. The *response epoch* follows the stimulus epoch, signaled by 𝑢*_fix_* being reset to 0. The network response and associated error were evaluated during the response epoch period. For the memoryless Go task, the target response must be equal to the active stimulus input (𝑢*_stim_*_1_ or 𝑢*_stim_*_2_) in that trial (which would be either +1 or –1). The duration of the response epoch randomly switched between 400 and 800 ms, again to ensure that the network learned the task and not the specific timings. Thus, the minimum trial duration was 800 ms (200+200+400 across the three epochs), while the maximum trial duration was 1800 ms (200+800+800). During training, the choice between 𝑢*_stim_*_1_and 𝑢*_stim_*_2_, the specific input (–1 or +1) to the chosen stimulus, and the durations of the stimulus and the response epochs were randomly varied across trials.

### Training and testing procedures

The input and recurrent synaptic weights were modified at the end of each trial based on a modified form of reward-modulated Hebbian learning (Miconi, 2017). During a trial, the potential weight change or eligibility trace for each recurrent synapse from the 𝑗^th^ unit to the 𝑖^th^ unit was accumulated over the course of the trial as:

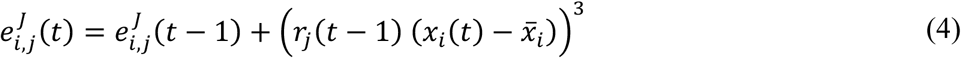

where 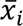 maintained a short-term running weighted average of the 𝑥*_i_* calculated as 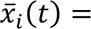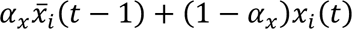. Thus, 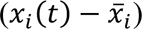 tracked sudden fluctuations in the unit’s activity, primarily due to the activity perturbation Δ*_i_*. Similarly, for each synapse from 𝑘^th^ external input to 𝑖^th^ unit, the eligibility trace 𝑒*_i_*_*i,k*_ was:

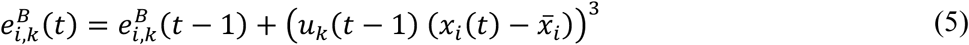

At the end of each trial, error was calculated over the response epoch (RE) as the average absolute difference between the network output (𝑟*_obs_*(𝑡)) and expected output (𝑟*_exp_*(𝑡)) for the trial:

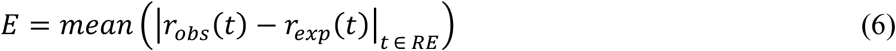

The reward for the trial was taken as the negative of the error, 𝑅 = −𝐸. The reward prediction error 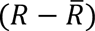 signal was then computed with 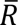, the expected reward, calculated for trial 𝑛 as:

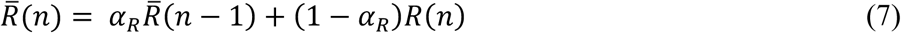

The reward prediction error was used to modulate the eligibility traces and compute weight changes:

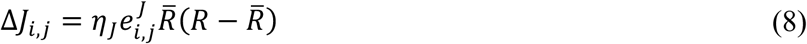

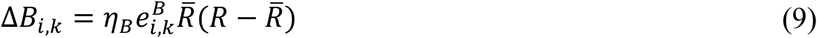

where 𝜂_5_and 𝜂_7_defined the learning rates for the ***J*** and ***B*** matrices, respectively. In these simulations, the default values of the relative weightages for computing 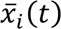 and 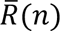 were 𝛼*_x_* = 0.05 and 𝛼*_R_* = 0.33, respectively. The default learning rate parameters were set at 𝜂_J_ = 𝜂_B_ = 0.15. Synaptic weights were updated using these relationships and the network then switched to the next trial of the cognitive task to repeat the whole updation process again. Under default conditions, convergence of the network (completion of training) was defined to be achieved if the error value 𝐸 was less than 0.05 for a threshold number of consecutive trials (𝑇_th_). The default value of 𝑇_AB_was taken as 100.

By default, trained networks were tested without the activity perturbation. Each network was tested on 16 different trials covering all 4 combinations of inputs (either 𝑢*_stim_*_1_ or 𝑢*_stim_*_2_ taking a value of either –1 or +1) and 4 trial durations (800, 1200, 1400, or 1800 ms). The network performance was then quantified based on the absolute error, 𝐸, computed for each trial. A trial was counted as a correct trial if 𝐸 < 0.05.

### Analyzing latent network dynamics

The response of all the units in the network across all trials were concatenated into a matrix *R* of dimensions 𝑇 × 𝑁 where 𝑁 defined the total number of recurrent units and 𝑇 represented the total number of time points across all 16 trials. To obtain the latent space dynamics of each network, we performed principal component analysis (PCA) on the associated 𝑅 matrix. The projections to the first three principal components of the activity of the 𝑁 recurrent units across time, during different kinds of trials, were used for visualizing the latent space trajectories in 3D. The dimensionality of the system was computed using the eigen values (λ*_i_*) as follows (Abbott et al., 2011; Mazzucato et al., 2016; Litwin-Kumar et al., 2017):

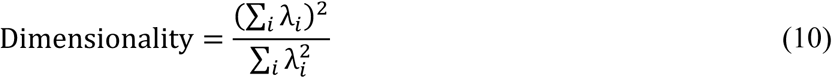

Since latent space dynamics of different networks trained on the same task form different trajectories in different latent spaces, the latent spaces themselves need to be aligned before the network trajectories can be compared across networks. Canonical correlation analysis (CCA) applies linear transformations to find a new set of dimensions for the latent space dynamics of a pair of networks, 𝐿_1_ and 𝐿_2_, such that their pairwise correlation is maximized (Gallego et al., 2020). When 𝑀_1_and 𝑀_2_ defined the coefficient matrices for projecting 𝐿_1_and 𝐿_2_ to the new dimensions, respectively, the projections of the latent space dynamics along the new dimensions were calculated as:

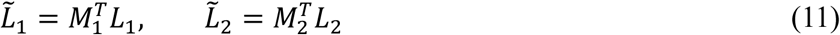

Then, the latent space dynamics of network 2 aligned with network 1 was obtained as:

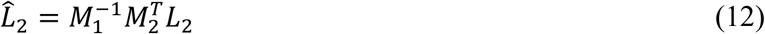

For each network, the projections along the first 4 PCs were taken for CCA transformation and the latent space trajectories of each pair of networks were aligned. The dimensionality value calculated using equation 10, across all 318 networks was around 3. As dimensionality approximately represents the number of dimensions required for explaining around 80% of the total variance, we used 4 dimensions for the CCA transformation. Therefore, the 𝑀*_i_* matrices were of dimension 4 × 4. For all our analyses, network N1 was used as the reference network for mappings.

To observe the variability in latent space dynamics due to change in network configurations, Euclidean distances between trajectories in the aligned latent space was computed for each pair of networks. Let 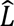 and 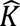 be 𝑇 × 𝑚 matrices of aligned latent space trajectories of the dynamics of two networks, where 𝑇 defined the number of time points and 𝑚 = 4 denoted the number of dimensions of the aligned trajectories. The distance between the trajectories as a function of time was then calculated as:

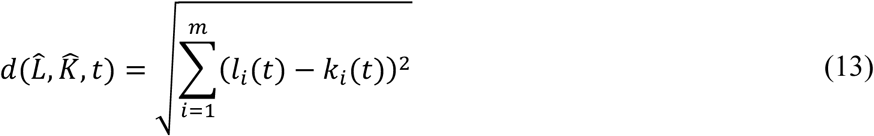

where 𝑙*_i_*(𝑡) and 𝑘*_i_*(𝑡) defined the columns of the matrix 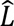 and 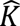 respectively. To quantify the impact of heterogeneity level on network dynamics, the distance between the heterogeneous (H1-H5) network trajectories were calculated with respect to the homogenous (H0) network trajectory.

### Assessing the robustness of networks to post-training introduction of heterogeneities

The different kinds of post-training heterogeneities for which robustness of all 318 networks was assessed are described below (Supplementary Table S1). Unless otherwise stated, the performance error and dynamics of the networks with post-training heterogeneities (P1–P5) were compared to that of the respective P0 network (where there was no post-training heterogeneity). In each of the cases below, to explore the robustness of the network against the post training heterogeneities, the difference in the absolute trial errors with and without heterogeneities were calculated. Similarly, the impact of post-training heterogeneities on the latent space dynamics was quantified as the distance of trajectories with post-training heterogeneity (P1–P5) from the respective P0 trajectory.

#### Intrinsic heterogeneity in unit time constants

We tested two types of post-training intrinsic heterogeneities in time constant distributions. First, the extent of heterogeneities in terms of the distance of time constants from the mean was kept constant, while the mean time constant was gradually shifted. The shifts in the time constant of each unit (in ms) were (0, 1, 5, 10, 20), in that sequence, for P0 to P5 levels of post-training heterogeneities. In this case, we used the respective P0 network as the reference for computing errors and distances (the default scenario mentioned above). In a second scenario, we kept the mean to be the same with the variance of post-training heterogeneities changing across P0–P5 (with P0 representing zero variance). As this was identical to how H0–H5 were defined, a network trained with Hx showed minimal error for Px. Therefore, in this case, errors and distances were computed with reference to the heterogeneity level that the specific network was trained with.

#### Jitter in recurrent synaptic weights

To assess the robustness of networks to changes in the trained values of recurrent synaptic weights, we introduced Gaussian jitter with zero mean and increasing variance. The standard deviation values of the Gaussian noise for P0–P5 were (0, 0.01, 0.02, 0.03, 0.04, 0.05).

#### Distribution pattern of initial activity

These post-training heterogeneities reflect differences in initial activity patterns at the beginning of the trials (𝒙(0)). We tested two types of post-training heterogeneities in initial activity patterns here. First, the extent of heterogeneities in terms of the initial activity patterns from the mean was kept constant, while the mean activity value was gradually shifted. The shifts in the initial activity value of each unit were (0, 0.1, 0.25, 0.5, 1, 2), in that sequence, for P0 to P5 levels of post-training heterogeneities. In a second scenario, we kept the mean activity to be the same with the range of post-training heterogeneities changing across P0–P5 (with P0 representing zero perturbations). The range of heterogeneities in the initial activity value of each unit were (0, 0.1, 0.25, 0.5, 1, 2), in that sequence, for P0 to P5 levels of post-training heterogeneities. For instance, for the P5 value, the range would be –2 to +2 for how the initial activity heterogeneities will be introduced. At P0, 𝒙(0) = 0.

#### Stimulus epoch duration

During the training process, we had used a stimulation epoch duration of either 200 ms or 800 ms. We tested the robustness of the learnt networks to changes in the stimulus epoch duration. The stimulus epoch durations for P0–P5 post-training heterogeneities were (800, 600, 400, 200, 100, 50) in that sequence.

#### Activity perturbation frequency

While activity perturbations were introduced during training process, they were turned off during testing. We tested the robustness of the learnt networks to introduction of increasing frequencies of activity perturbation. The frequency of activity perturbations (in Hz) for P0–P5 post-training heterogeneities, in that sequence, were (0, 2, 4, 6, 8, 10).

### Computational details

All simulations and analyses were performed within the MATLAB (Mathworks Inc.) programming environment.

## ACKNOWLEDGMENTS

The authors thank Dr. Alex Cayco-Gajic, Dr. Laura Driscoll, and Dr. Sara Solla for helpful discussions. The authors thank members of the cellular neurophysiology laboratory for helpful discussions and for comments on a draft of this manuscript. This work was supported by the Wellcome Trust-DBT India Alliance (Senior fellowship to R. N.; IA/S/16/2/502727), Pratiksha Trust (A.S.), and the Ministry of education (A. S. and R. N.).

## Author contributions

A. S. and R. N. designed experiments; A. S. performed experiments; A. S. analyzed data; A. S. and R. N. wrote the paper.

