## Supplementary Figures S1-S6 and Supplementary Table S1 for "Diversity in the impact of heterogeneities on recurrent networks performing a cognitive task"

### Supplementary material

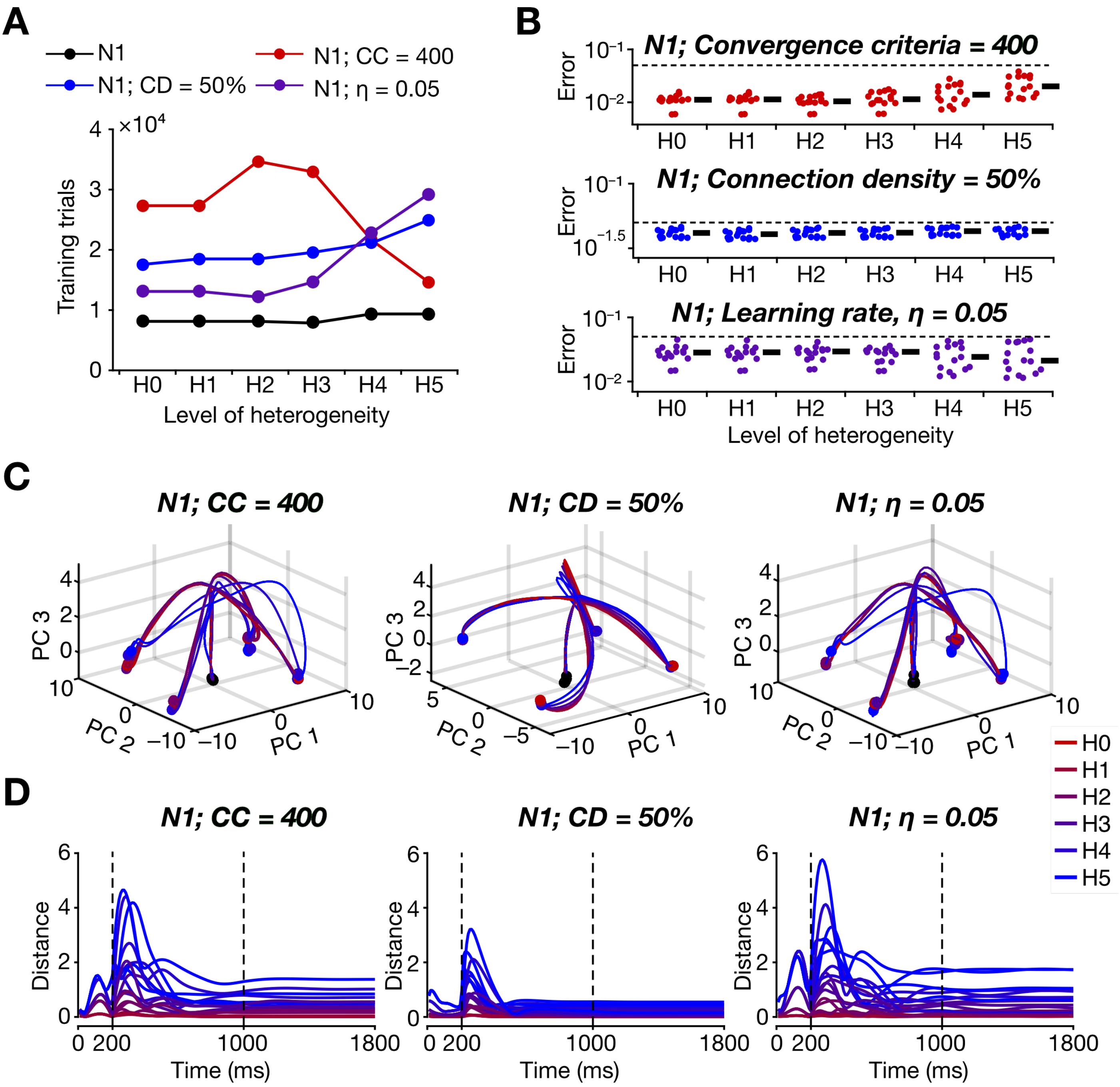

**Supplementary Figure S1. Training performance and dynamics of heterogeneous recurrent networks with more stringent hyperparameters.** *A:* Number of trials taken to train network N1 to reach convergence criteria with default hyperparameters (N1), with more stringent convergence criteria (N1; CC=400), with reduced connection probability between neurons (N1; CD=50%), and reduced learning rate (N1; $\eta$=0.05). For the default network (N1), the convergence criterion required error to be below threshold for 100 trials, the connection probability was 100%, and learning rate $\eta$=0.15. *B:* Bee-swarm plot of distribution of errors for different levels of training heterogeneity with each of the different groups. The black dashed line represents the threshold error (0.05) below which the trial is considered a correct trial. Solid lines represent the median values for each heterogeneity level. *C:* CCA-aligned trajectories of networks with the different stringent training criteria, with all 6 levels of heterogeneities (H0–H5). *D:* The distance between latent-space trajectories for all levels of heterogeneity with respect to their respective H0 network shown for all training heterogeneity levels.

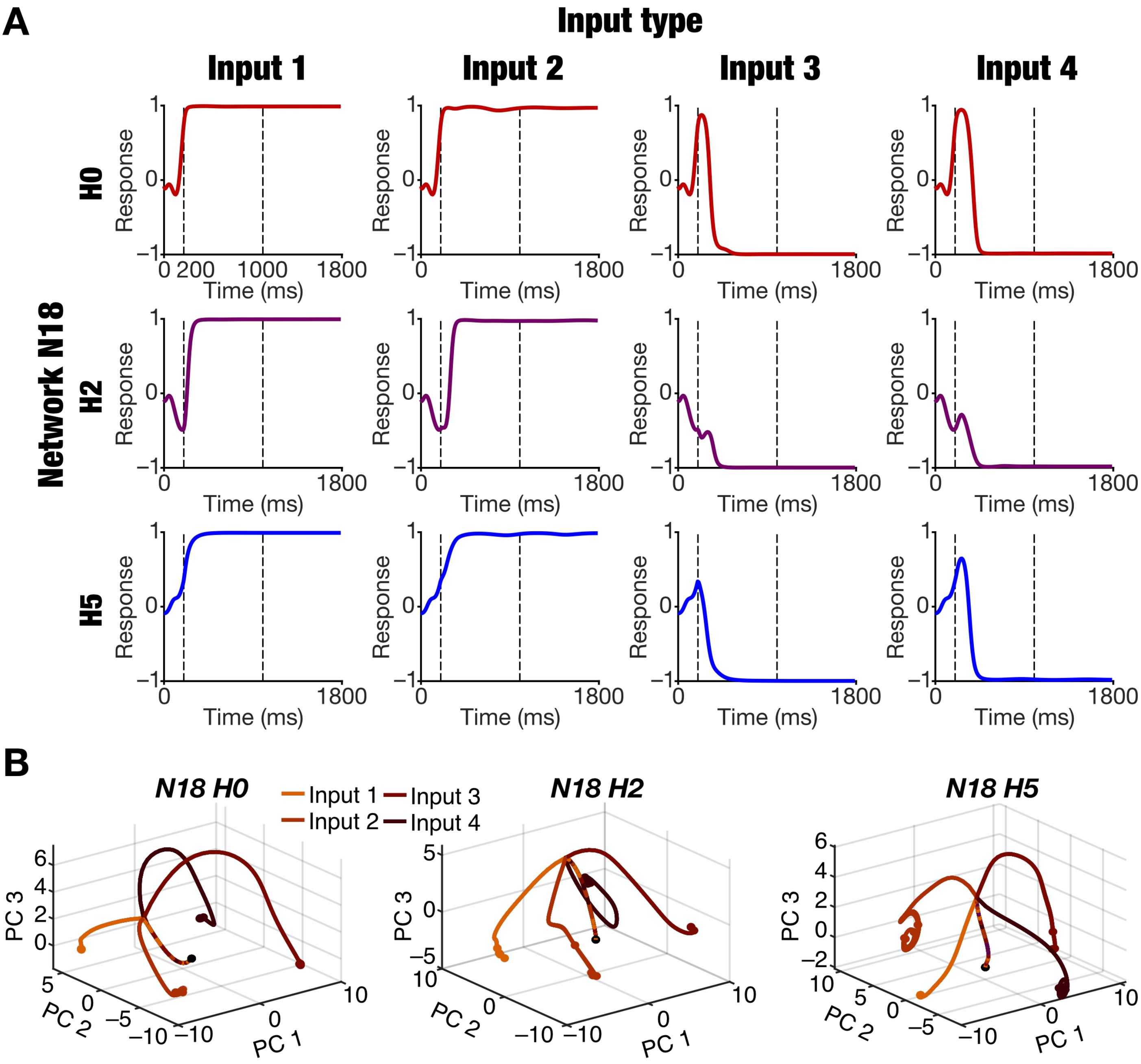

**Supplementary Figure S2. Dynamics of recurrent networks with the same hyperparameters and different levels of training heterogeneities.** *A:* Temporal evolution of the response of the output unit in network N18 trained with three different levels of heterogeneities (H0, H2, and H5). Shown are responses of the output unit to the four distinct input types (**Input 1**: $u_{stim1} = 1,\ u_{stim2} = 0$; **Input 2**: $u_{stim1} = 0,\ u_{stim2} = 1$; **Input 3**: $u_{stim1} = -1,\ u_{stim2} = 0$ **Input 4**: $u_{stim1} = 0,\ u_{stim2} = -1$). It may be noted the output unit has converged to its respective expected output values in each of the four input types across all levels of heterogeneities. The variability in the temporal dynamics across different levels of heterogeneities, especially during the fixation epoch and the initial part of the stimulus epoch, may also be noted. *B:* The dynamics of all units in the network are represented by input-dependent trajectories in their respective reduced dimensional spaces, computed for each network using principal component analysis (PCA). These are shown for the same networks that are depicted in panel A. The input-dependence of the trajectories and the fixed points may be noted. Variability in the locations of fixed points and in the trajectories across the different levels of heterogeneities may also be noted.

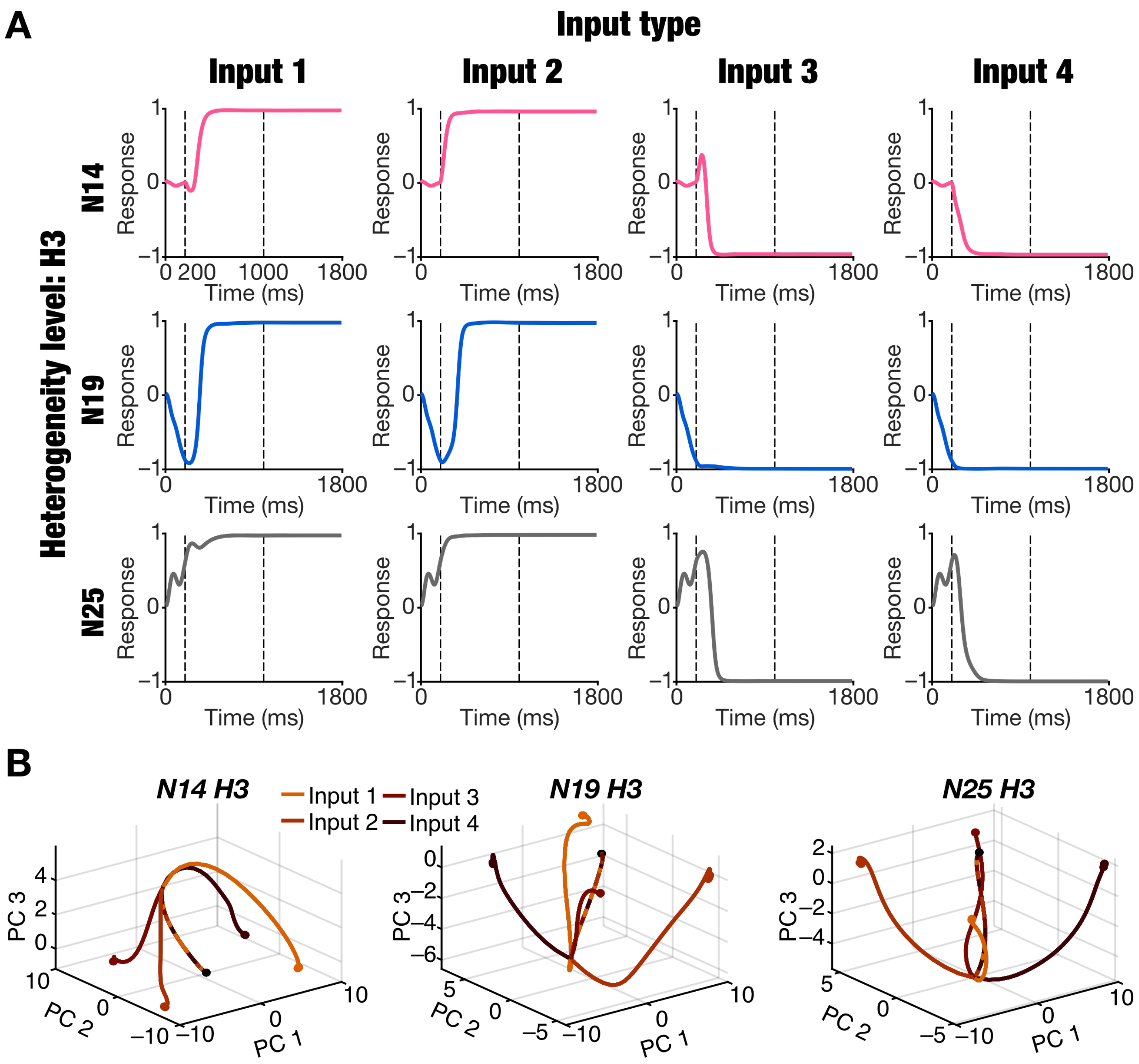

**Supplementary Figure S3. Dynamics of recurrent networks with distinct hyperparameters and same level of training heterogeneities.** *A:* Temporal evolution of the response of the output unit in three different networks (N14, N19, and N25) trained with the same levels of heterogeneity (H3). Shown are responses of the output unit to the four distinct input types (**Input 1**: $u_{stim1} = 1$, $u_{stim2} = 0$; **Input 2**: $u_{stim1} = 0$, $u_{stim2} = 1$; **Input 3**: $u_{stim1} = -1$, $u_{stim2} = 0$ **Input 4**: $u_{stim1} = 0$, $u_{stim2} = -1$). It may be noted the output unit has converged to its respective expected output values in each of the four input types across all networks. The variability in the temporal dynamics across different networks, especially during the fixation epoch and the initial part of the stimulus epoch, may also be noted. *B:* The dynamics of all units in the network are represented by input-dependent trajectories in their respective reduced dimensional spaces, computed for each network using principal component analysis (PCA). These are shown for the same networks that are depicted in panel A. The input-dependence of the trajectories and the fixed points may be noted. Variability in the locations of fixed points and in the trajectories across the different networks may also be noted.

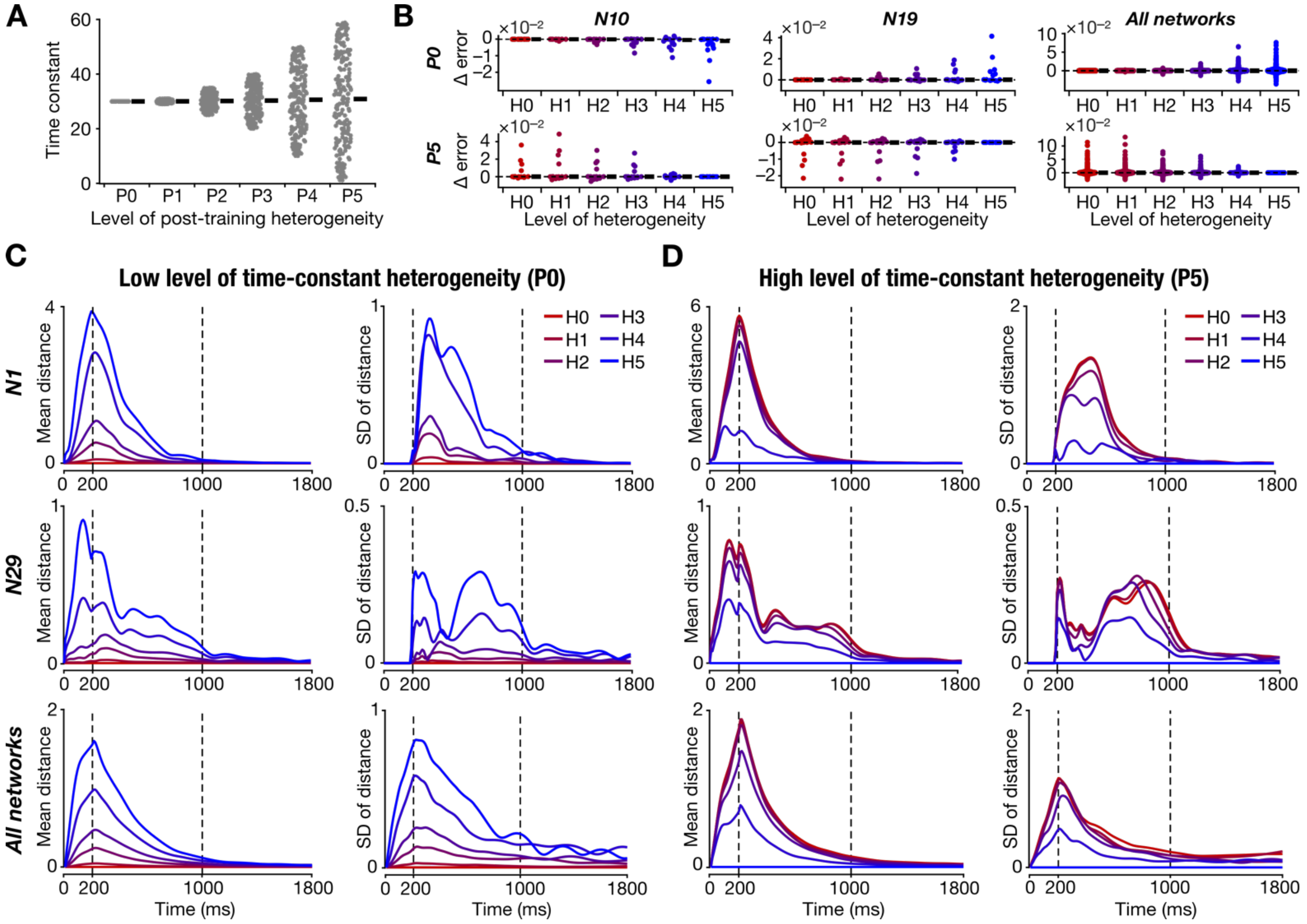

**Supplementary Figure S4. Pronounced network-to-network variability in the robustness of heterogeneous recurrent networks to post-training intrinsic heterogeneities.** *A:* Distribution of time constants for six different levels of post-training heterogeneity (P0–P5). Note that the mean (black bar) remained the same, while the range of time constants increased as the level of post-training heterogeneities are increased. This is akin to heterogeneities introduced during training (H0–H5; Fig. 1) and constitutes a shift in level of heterogeneity post-training. Therefore, errors and distances in panels B–D were calculated with respect to the heterogeneity level the networks were trained with. For instance, when networks trained with H0–H5 are subjected to perturbation P5, the outcomes are compared with H5 network. *B:* Distribution of the difference in response errors (across different trials) for low (P1; top row) and high (P5; bottom row) levels of post-training heterogeneity, introduced in networks trained with H0–H5 heterogeneities. The first column shows an example network (N10) that showed a reduction in error with increase in level of training heterogeneities (H0–H5). The second column depicts another network (N19) where the error increased in networks trained with higher level of heterogeneities. The third column summarizes the difference in errors across all 53 networks, spanning different trials. Black bars represent the median values. *C:* Mean (left column) and standard deviation (SD; right column) of the distance distribution (computed from trajectories in the latent space) with respect to H0 for low level of post-training heterogeneity (P0). Each panel shows plots for networks trained with different levels of heterogeneity (H0–H5). The first row shows an example network (N1) where the mean distance and the associated variability were high. The second row depicts another network (N29) where the mean distance and the associated variability were low. The third row depicts mean distance and the associated variability across all 53 networks. *D:* Same as *C* but for P5 instead of P0. For panels C and D, the error for the trained level of heterogeneity is low and increases progressively with increasing distance from the trained level of heterogeneity.

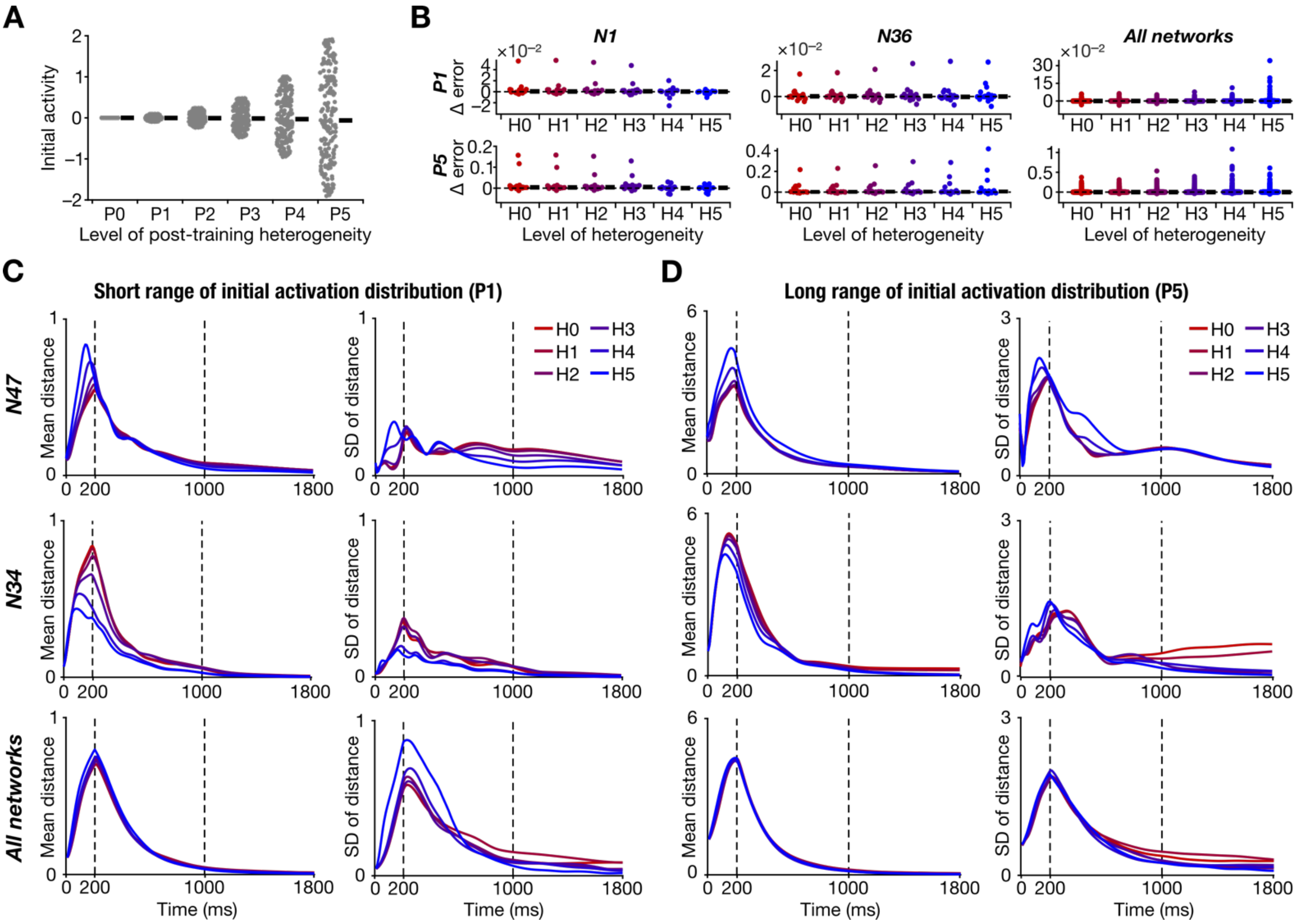

**Supplementary Figure S5. Pronounced network-to-network variability in the robustness of heterogeneous recurrent networks to post-training shift in network activity at task initiation.** *A:* Distribution of network activity range in all recurrent units at network initiation for six different levels of post-training heterogeneity (P0–P5). Note the shift in the range of the initial activity distribution (without shift in mean) with increasing in level of post-training heterogeneity. *B:* Distribution of the difference in response errors (across different trials) with respect to P0 for low (P1; top row) and high (P5; bottom row) levels of post-training heterogeneity, introduced in networks trained with H0–H5 heterogeneities. The first column shows an example network (N1) that showed a reduction in error with increase in level of training heterogeneities (H0–H5). The second column depicts another network (N36) where the error was high in networks trained with higher level of heterogeneities. The third column summarizes the difference in errors across all 53 networks, spanning different trials. Black bars represent the median values. *C:* Mean (left column) and standard deviation (SD; right column) of the distance distribution (computed from trajectories in the latent space) with respect to P0 for low level of post-training heterogeneity (P1). Each panel shows plots for networks trained with different levels of training heterogeneity (H0–H5). The first row shows an example network (N47) where the mean distance and the associated variability increased with increasing level of training heterogeneities. The second row depicts another network (N34) where the mean distance and the associated variability decreased with higher level of heterogeneities introduced during training. The third row depicts mean distance and the associated variability across all 53 networks. *D:* Same as *C* but for P5 instead of P1.

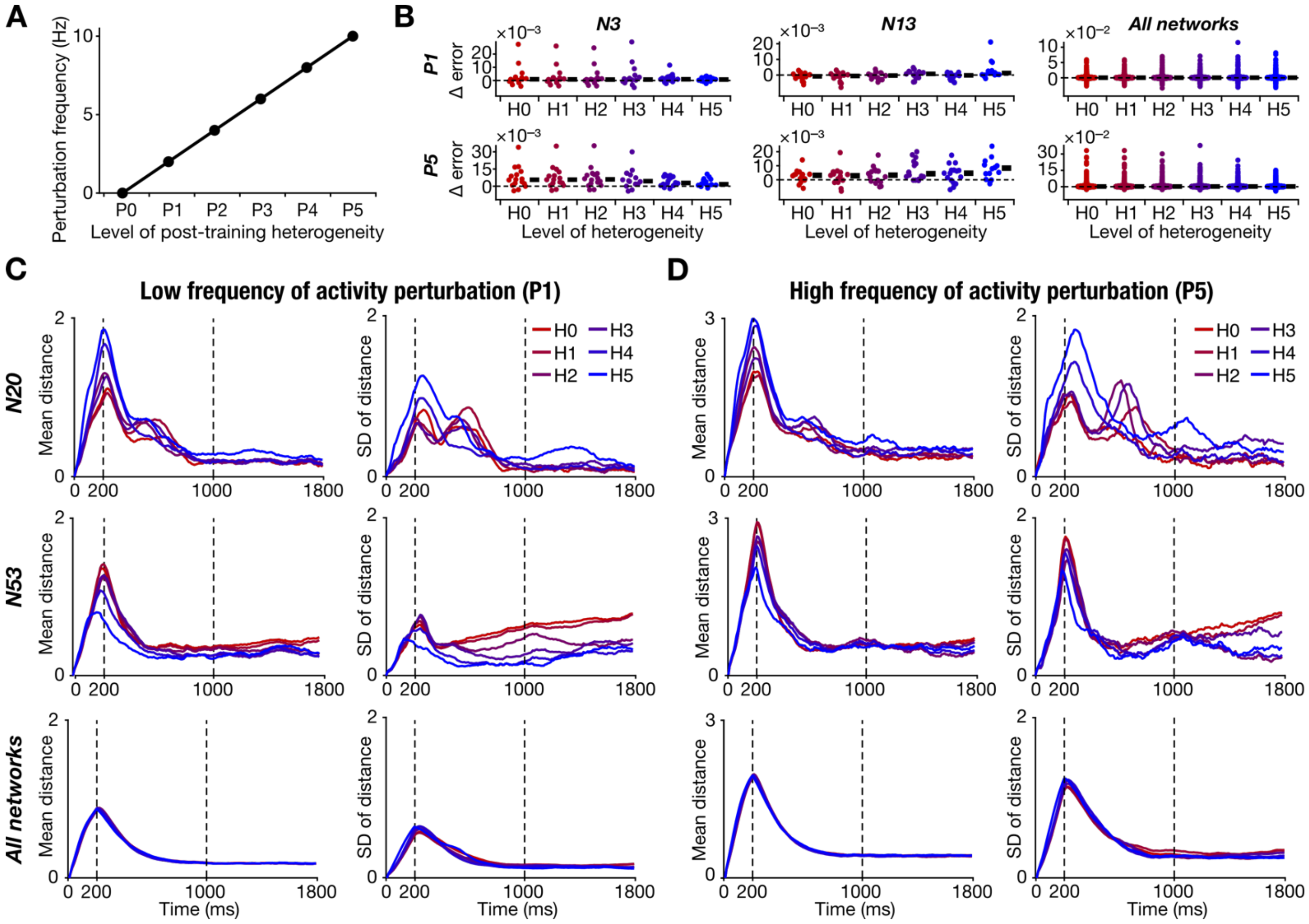

**Supplementary Figure S6. Pronounced network-to-network variability in the robustness of heterogeneous recurrent networks to post-training activity perturbation.** *A:* Increasing frequency of activity perturbation introduced for six different levels of post-training heterogeneity (P0–P5). *B:* Distribution of the difference in response errors (across different trials) with respect to P0 for low (P1; top row) and high (P5; bottom row) levels of post-training heterogeneity, introduced in networks trained with H0–H5 heterogeneities. The first column shows an example network (N3) that showed a reduction in error with increase in level of training heterogeneities (H0–H5). The second column depicts another network (N13) where the error was high in networks trained with higher level of heterogeneities. The third column summarizes the difference in errors across all 53 networks, spanning different trials. Black bars represent the median values. *C:* Mean (left column) and standard deviation (SD; right column) of the distance distribution (computed from trajectories in the latent space) with respect to P0 for low level of post-training heterogeneity (P1). Each panel shows plots for networks trained with different levels of training heterogeneity (H0–H5). The first row shows an example network (N20) where the mean distance and the associated variability increased with increasing level of training heterogeneities. The second row depicts another network (N53) where the mean distance and the associated variability decreased with higher level of heterogeneities introduced during training. It may be noted that the variability in distances were high for the N53 network even during the response epoch, especially when trained with low level of heterogeneities (H0–H1). The third row depicts mean distance and the associated variability across all 53 networks. *D:* Same as *C* but for P5 instead of P1.

**Supplementary Table S1. Details of the different levels of the various post-training heterogeneities.**

| Type of post–training heterogeneity | Description of heterogeneity | Level of Heterogeneity | Values |
|---|---|---|---|
| **Mean of time constant distribution (Fig. 5)** | Time constants of each unit, $\tau$, are shifted by $\tau_{shift}$ such that the new time constants are $\tau + \tau_{shift}$ | P0 | $\tau_{shift} = 0$ |
| | | P1 | $\tau_{shift} = 1$ |
| | | P2 | $\tau_{shift} = 5$ |
| | | P3 | $\tau_{shift} = 10$ |
| | | P4 | $\tau_{shift} = 20$ |
| | | P5 | $\tau_{shift} = 40$ |
| **Range of time constant distribution (Supplementary Fig. S4)** | $\tau$ is sampled from a uniform distribution spanning $\tau_{range}$ with the same seed value used for training | P0 | $\tau_{range} = 30$ |
| | | P1 | $\tau_{range} = [29,31]$ |
| | | P2 | $\tau_{range} = [25,35]$ |
| | | P3 | $\tau_{range} = [20,40]$ |
| | | P4 | $\tau_{range} = [10,50]$ |
| | | P5 | $\tau_{range} = [1,59]$ |
| **Noise in recurrent synaptic weight (Fig. 6)** | Noise $\boldsymbol{J}^{noise}$ sampled from a normal distribution $\mathcal{N}(0, \sigma^2)$ were added to the recurrent synaptic weights, $\boldsymbol{J}$ | P0 | $\sigma = 0$ |
| | | P1 | $\sigma = 0.01$ |
| | | P2 | $\sigma = 0.02$ |
| | | P3 | $\sigma = 0.03$ |
| | | P4 | $\sigma = 0.04$ |
| | | P5 | $\sigma = 0.05$ |
| **Initial activity distribution mean (Fig. 7)** | $x(0)$ is sampled from a uniform distribution of range $x_{shift} + [-0.1,0.1]$ | P0 | $x_{shift} = 0$ |
| | | P1 | $x_{shift} = 0.1$ |
| | | P2 | $x_{shift} = 0.25$ |
| | | P3 | $x_{shift} = 0.5$ |
| | | P4 | $x_{shift} = 1$ |
| | | P5 | $x_{shift} = 2$ |
| **Initial activity distribution range (Supplementary Fig. S5)** | $x(0)$ is sampled from a uniform distribution of range $[-x_{range}, x_{range}]$ | P0 | $x_{range} = 0$ |
| | | P1 | $x_{range} = 0.1$ |
| | | P2 | $x_{range} = 0.25$ |
| | | P3 | $x_{range} = 0.5$ |
| | | P4 | $x_{range} = 1$ |
| | | P5 | $x_{range} = 2$ |
| **Stimulus epoch duration (Fig. 8)** | The duration of the stimulus epoch was set to $T_{SE}$ | P0 | $T_{SE} = 800$ |
| | | P1 | $T_{SE} = 600$ |
| | | P2 | $T_{SE} = 400$ |
| | | P3 | $T_{SE} = 200$ |
| | | P4 | $T_{SE} = 100$ |
| | | P5 | $T_{SE} = 50$ |
| **Activity perturbation frequency (Supplementary Fig. S6)** | Activity perturbations, $\Delta$, are introduced to each unit with the frequency $f_\Delta$ | P0 | $f_\Delta = 0$ |
| | | P1 | $f_\Delta = 2$ |
| | | P2 | $f_\Delta = 4$ |
| | | P3 | $f_\Delta = 6$ |
| | | P4 | $f_\Delta = 8$ |
| | | P5 | $f_\Delta = 10$ |
